# Mechanisms of HSV-1 helicase–primase inhibition and replication fork complex assembly

**DOI:** 10.64898/2025.12.23.696259

**Authors:** Zishuo Yu, Pradeep Sathyanarayana, Cong Liu, Pan Yang, Sandra K Weller, Mrinal Shekhar, Donald M. Coen, Joseph J. Loparo, Jonathan Abraham

**Affiliations:** Department of Microbiology, Blavatnik Institute, Harvard Medical School, Boston, MA, USA; Department of Biological Chemistry and Molecular Pharmacology, Blavatnik Institute, Harvard Medical School, Boston, MA, USA; Center for the Development of Therapeutics, Broad Institute of Harvard and MIT, Cambridge, MA, USA; Department of Molecular Biology and Biophysics, University of Connecticut School of Medicine, Farmington, Connecticut, USA; Department of Medicine, Division of Infectious Diseases, Brigham and Women’s Hospital, Boston, MA, USA; Center for Integrated Solutions for Infectious Diseases, Broad Institute of Harvard and MIT, Cambridge, MA, USA; Howard Hughes Medical Institute, Boston, MA, USA

**Keywords:** Herpesvirus, helicase, primase, polymerase, antiviral

## Abstract

Herpesviruses are widespread double-stranded DNA viruses that establish lifelong latency and cause various diseases. Although DNA polymerase-targeting antivirals are effective, increasing drug resistance underscores the need for alternatives. Helicase–primase inhibitors (HPIs) are promising antivirals, but their mechanisms of action are poorly defined. Furthermore, how the helicase–primase (H/P) complex and DNA polymerase coordinate genome replication is not well understood for herpesviruses. Here, we report cryo-EM structures of the herpes simplex virus (HSV) H/P complex bound to HPIs, showing that these lock the helicase-primase complex in an inactive conformation. Single-molecule assays reveal that HPIs cause helicase-primase complexes to pause in unwinding activity on DNA. The structure of an HPI-bound replication fork complex, comprising the H/P complex (UL5, UL52, and UL8) and polymerase holoenzyme (UL30 and UL42), reveals a previously uncharacterized interface bridging these complexes. These findings provide a structural framework for understanding herpesvirus replisome assembly and advancing inhibitor development.

## Introduction

Herpesviruses are a large group of enveloped double-stranded DNA (dsDNA) viruses that contain many highly prevalent human pathogens.^1^ There are three subfamilies of herpesviruses—α, β, and γ.^1^ Herpes simplex virus (HSV) belongs to the α-herpesvirus subfamily, which is characterized by rapid replication kinetics and a broad host range, with many members establishing lifelong latency in sensory neurons. There are two HSV serotypes: HSV-1 is primarily associated with orolabial infections, and HSV-2 is primarily associated with genital ulcers.^2^ HSV can cause meningoencephalitis in immunocompetent individuals, and severe, disseminated disease in immunocompromised individuals.^2^ The β-herpesvirus subfamily includes human cytomegalovirus (HCMV), which is associated with congenital birth defects and severe disease in immunocompromised individuals.^3^ The γ-herpesvirus subfamily includes Epstein–Barr virus (EBV) and Kaposi’s sarcoma-associated herpesvirus (KSHV), which are oncogenic and respectively cause lymphomas and nasopharyngeal carcinoma, and Kaposi’s sarcoma.^4,5^

Five HSV proteins form the core replication machinery at the DNA replication fork. These are the heterotrimeric helicase–primase (H/P), which comprises UL5 (helicase), UL52 (primase), and UL8 (accessory protein), and the heterodimeric polymerase holoenzyme, which comprises a catalytic subunit (UL30) and a processivity factor (UL42).^6^ The HSV H/P complex has 5′ to 3′ helicase, ATPase, primase, and DNA-binding activities.^7–9^ UL5 belongs to the Superfamily 1 (SF1) helicases, which is a diverse group of monomeric and dimeric helicases.^10–12^ UL52 is a member of the archaeo-eukaryotic primase (AEP) superfamily, with a catalytic core (AEP-like domain).^13^ Upon recruitment to the viral replication forks, UL5 further unwinds the dsDNA on the leading strand, while UL52 synthesizes short RNA primers on the lagging strand to provide substrates for DNA synthesis.^6^ UL8 is an accessory protein required for nuclear localization of the H/P complex and aids in unwinding dsDNA.^14^ Based on bioinformatics analysis, UL8 is predicted to resemble a catalytically dead family B DNA polymerase, with palm, thumb, and fingers domains.^15^ A sixth protein, the single-strand DNA (ssDNA)-binding protein, infected cell protein 8 (ICP8, UL29), is an important component of the replication machinery.^6^

The HSV polymerase is a B family DNA polymerase with 3′–5′ proofreading activity.^16^ UL30 contains thumb, palm, fingers, a 3′–5′ exonuclease domain, and an N-terminal domain (NTD).^17^ The NTD contains a pre-NH_2_ terminal domain (preN) domain specific to herpesviruses that is important for HSV-1 replication but whose function is unclear.^17–19^ The processivity factor UL42 enhances DNA polymerase affinity for viral DNA and facilitates long-chain DNA synthesis.^20–22^ UL42 shares the same fold as eukaryotic proliferating cell nuclear antigen (PCNA), which forms rings around DNA.^23^ However, UL42 functions as a monomer without encircling the DNA.^24–27^

Among herpesviruses, approved antivirals are only available against HSV, varicella-zoster virus (VZV, another α-herpesvirus), and HCMV (a β-herpesvirus).^28^ Most current drugs against HSV target the viral DNA polymerase. Acyclovir and its orally available prodrug valacyclovir, which are nucleoside analogs that require activation by the viral thymidine kinase, have been used to treat HSV infections for several decades.^29^ However, these drugs are only partially effective at preventing virus reactivation.^30^ Foscarnet, a pyrophosphate analog that is a second-line drug, is associated with significant side effects. Importantly, mutations altering the viral thymidine kinase that actives acyclovir, and mutations in the catalytic subunit of the polymerase, UL30, can cause clinically relevant resistance to acyclovir and foscarnet.^31^ The emergence of drug-resistant HSV strains, particularly in immunosuppressed individuals, has made the identification of alternative antiviral strategies a priority.

The HSV H/P complex is a promising target to meet the need for additional antivirals against herpesviruses. Helicase-primase inhibitors (HPIs) that are under investigation for treating HSV are pritelivir and the pritelivir-derived agent IM-250, which are in phase III and I clinical trials, respectively.^32–35^ Amenamevir is an HPI that is approved for clinical use in Japan against HSV and VZV.^36,37^ How HPIs interfere with helicase-primase activity is unknown. Additionally, current HPIs have limited antiviral spectra of activity, and a lack of knowledge about their binding sites and mechanisms of action has hindered the development of HPIs for other herpesviruses.

Here, we used single-particle cryo-electron microscopy (cryo-EM) to obtain structures of DNA- and HPI-bound HSV helicase-primase complexes. The structures, paired with single-molecule experiments, reveal that HPIs lock the helicase-primase complex in an inactive conformation, causing it to pause in unwinding activity on DNA. The structure of an assembled replication fork complex also reveals an extensive H/P–polymerase interface, including a feature that is conserved among herpesviruses.

## Results

### HSV H/P complex structure determination

For structural studies, we purified the HSV H/P complex (UL5, residues 1–882; UL52, residues 1–1058; and UL8, residues 1–750) in transiently transfected mammalian cells (Figures 1a and S1a). When examined by mass photometry (MP), HSV H/P complex samples had a substantial proportion of particles with masses consistent with UL5/UL52/UL8 heterotrimers, which would have an expected mass of 293 kilodaltons (kDa) (Figure 1b and Table S1). We also observed peaks likely representing a UL5/UL52 subcomplex (202 kDa) and UL5 alone (92 kDa), suggesting that at the low concentration used for the MP experiments, there was some spontaneous dissociation of the complex.

**Figure 1.**
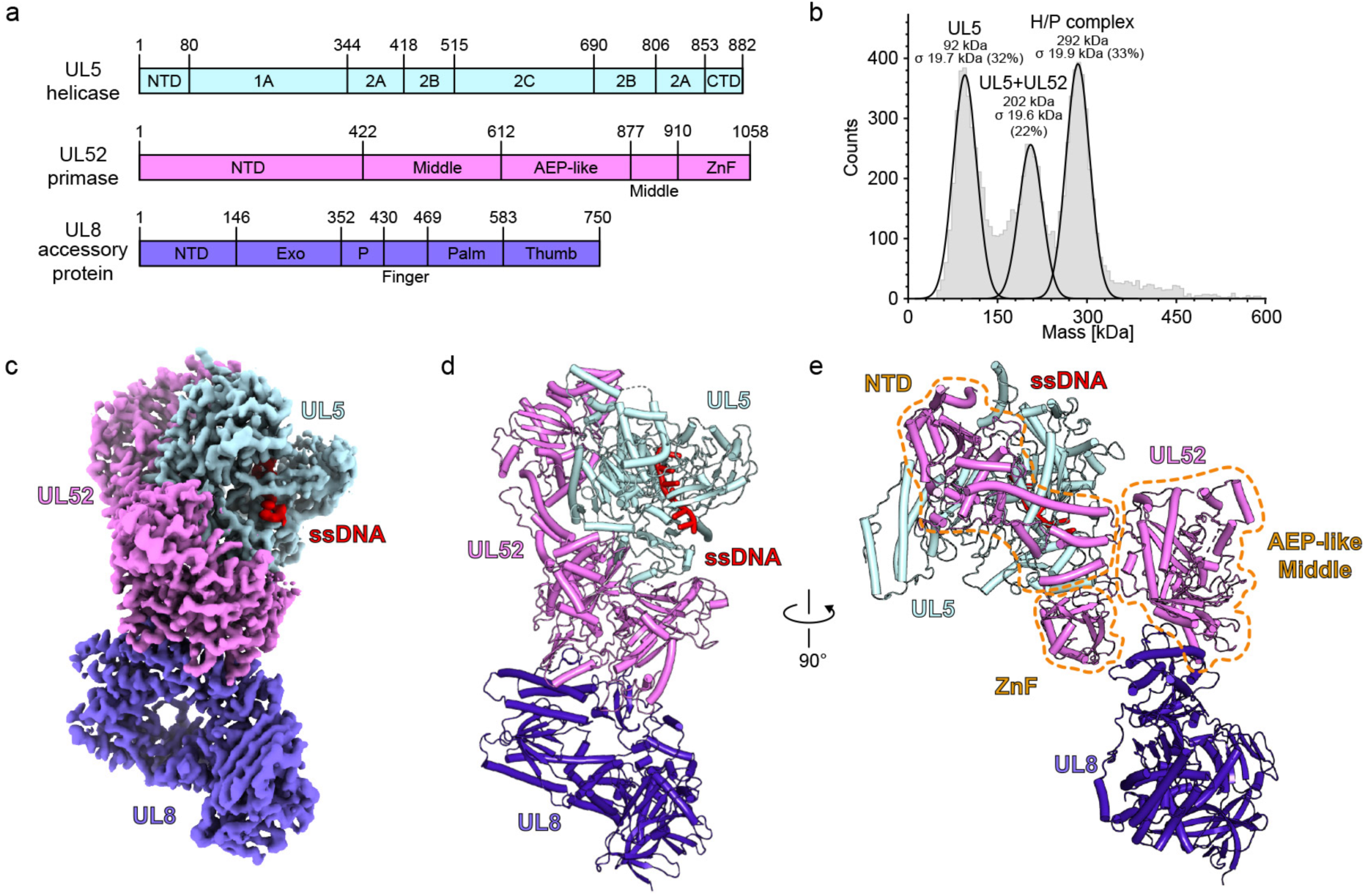
Overall structure of the HSV-1 H/P complex. (a) Domain architecture of components of the HSV H/P complex, which includes UL5 (helicase), UL52 (primase), and UL8 (accessory protein). NTD, N-terminal domain; CTD, C-terminal domain; AEP-like, archaeo-eukaryotic primase-like domain; ZnF, zinc finger domain; Exo, exonuclease domain; P, palm domain. (b) Mass photometry analysis of the HSV H/P complex. The experiment was performed twice, and representative data are shown. See Table S1 for additional information. (c) Cryo-EM map of the HSV H/P complex bound to ssDNA, colored as indicated. Only six nucleotides of ssDNA could be resolved in the map. (d and e) Ribbon diagram of the HSV H/P complex. Two views are shown. In panel E, the UL52 NTD, middle, AEP-like, and ZnF domains are circled.

We assembled the H/P complex onto an annealed forked DNA substrate in the presence of pritelivir and imaged samples by single-particle cryo-EM (Figure S1b and Table S2). We obtained a map of the complex bound to a 6-nucleotide ssDNA segment and pritelivir with a global resolution of 3.7 Å (Figures 1c, S1c and Table S3). We used focused refinement to obtain higher-resolution maps of specific regions of interest, with maps of UL5–UL52 interface at 3.4 Å and of the UL52–UL8 interface at 3.3 Å (Figures S1d and S1e).

### H/P complex architecture

The HSV H/P complex has a bilobed architecture centered on the UL52 primase, with UL5 and UL8 positioned on opposite sides of the complex (Figures 1c–1e). UL52 engages UL5 and UL8 through multiple structured subdomains (Figures 1d and S2). UL52 contains an NTD, a middle domain, an AEP-like domain (responsible for generating RNA primers), and a C-terminal zinc finger (ZnF) domain (Figure 1e). The UL52 NTD contacts the UL5 NTD, 1A, and CTD domains (Figure S2c). The UL52 ZnF contacts the UL5 2A domain (Figure S2d), and the UL52 middle domain makes extensive contacts with the UL8 thumb domain (Figure S2e). Notably, the UL52 AEP-like and ZnF domains are positioned near the exit of the helicase ssDNA channel (Figures S3a and S3b). This positioning would allow UL52 to initiate RNA primer synthesis on the DNA template as it emerges from the helicase.

### UL5 DNA recognition and ATP binding

The HSV UL5 helicase contains six domains: an NTD, four central domains—1A, 2A, 2B, and 2C—and a CTD (Figure 2a). UL5 domains 1A and 2A have the RecA-like fold characteristic of SF1 and SF2 helicases, while domain 2B has an SH3-like fold.^12^ The 2C domain, which folds as a bundle of ⍺-helices and is an insert in the 2B domain, is unique to herpesviruses and is absent in other SF1 helicases, leaving its function poorly understood. A similar 2C domain is in the *Saccharomyces cerevisiae* Pif1 DNA helicase, where it is hypothesized to interact with other proteins in cells^8,38^.

**Figure 2.**
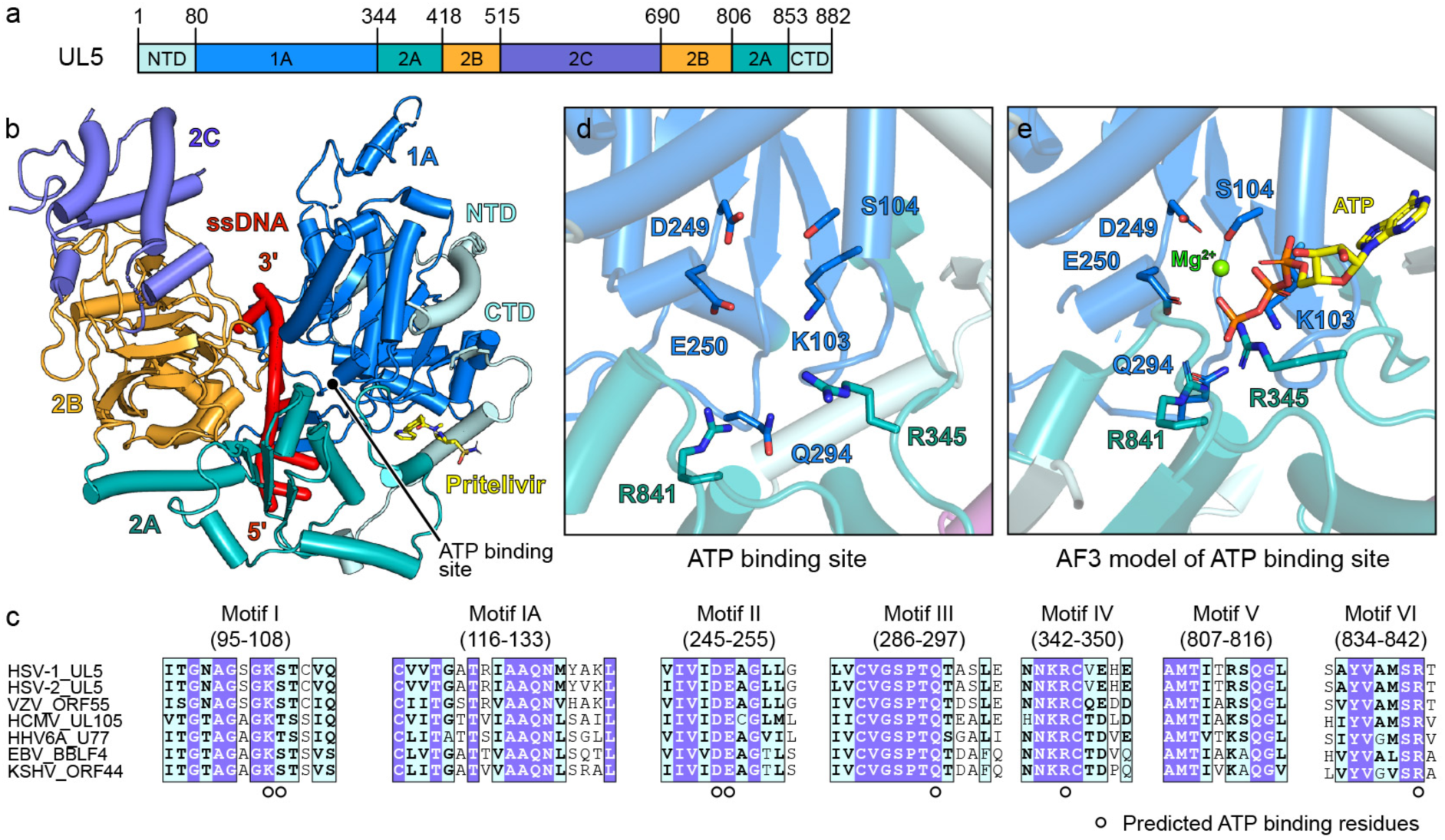
HSV UL5 helicase domains and ATP binding site. (a) HSV UL5 domain architecture. Domains 1A and 2A are RecA domains, domain 2B is an SH3-like fold domain, and domain 2C is a herpesvirus-specific domain. NTD, N-terminal domain; CTD, C-terminal domain. (b) HSV helicase UL5 bound to ssDNA. Six nucleotides of ssDNA could be resolved in the structure. The complex also included pritelivir, shown as sticks. (c) Sequence alignment of the UL5 active site. Conserved active site residues predicted to bind ATP are indicated. HSV-1, herpes simplex virus 1; HSV-2, herpes simplex virus 2; VZV, varicella-zoster virus; HCMV, human cytomegalovirus; EBV, Epstein–Barr virus; HHV-6A, human herpesvirus 6A; KSHV, Kaposi’s sarcoma-associated herpesvirus. (d and e) View of the ATP binding site in HSV UL5 in the cryo-EM structure of the H/P complex (d) or in an AlphaFold 3^40^ model of ATP-bound H/P complex (e). Conserved ATP-interacting residues are shown as sticks.

In the cryo-EM structure, UL5 binds single-stranded DNA (ssDNA) in a configuration consistent with other SF1 helicases.^12^ The UL5 2A domain is on the 5′ side of the ssDNA, and the UL5 1A domain is on the 3′ side. The ssDNA travels through a groove formed by domains 2A and 1A, flanked laterally by the 2B and 2C domains (Figure 2b).

UL5 contains seven conserved motifs common to the other SF1 helicases, which coordinate ATP binding, hydrolysis, and DNA translocation (Figure 2c). UL5 belongs to the SF1B subgroup, which translocates along ssDNA in the 5′–3′ direction. Upon ATP binding, conformational changes drive closure of the 2A/2B domains toward the 1A domain, promoting directional movement. Mutations in these conserved motifs abolish helicase activity and impair viral replication.^12,39^

Structural analysis shows that the seven conserved motifs, which form the ATP binding site, are located within the cleft between domains 1A and 2A (Figure 2d). Based on an AlphaFold 3^40^-predicted model of the ATP-bound H/P complex (Figures 2e and S4a), the key basic residues within the ATP binding site are K103 (motif I), R345 (motif IV), and R841 (motif VI), which coordinate ATP phosphate group binding, while S104 (motif I), D249 and E250 (motif II) coordinate a magnesium ion. The conserved residue Q294 (motif III) would be positioned to contact the γ-phosphate of ATP. Of the aforementioned residues, K103, D249, E250, and R345 have previously been demonstrated to be critical for helicase activity (Table S4).^10,41^ Overall, HSV UL5 shares a high degree of structural similarity with other SF1 helicases, particularly in its DNA recognition and ATP binding modes (e.g., as observed in the structure of ATP-bound *Sc*Pif1).^12,38^

### UL52 primase active site

Within the AEP core primase active site of UL52, three acidic residues that are conserved in herpesviruses—D628, D630, and D812—are positioned to coordinate a catalytic divalent metal ion required for the nucleotidyl transfer reaction (Figures S3c and S3e). Nearby, another conserved residue, R763, likely contributes to binding the phosphates of nucleotides. Supporting the structural analysis, the UL52 D628A and D630A mutations, which would disrupt metal ion coordination in the primase active site, have previously been demonstrated to decrease primase activity (Table S4).^13,42^ The ZnF domain, which is at the C-terminus of UL52, contains a coordinating center with four conserved residues—H993, C988, C1023, and C1028 (Figures S3d and S3e). The C1023A and C1028A ZnF domain mutations, which would impair zinc ion coordination, have previously been shown to cause UL5–UL52 subcomplexes to lose nearly all helicase and primase activities (Table S4).^43,44^

### A conserved H/P inhibitor binding mode

To determine the structural basis of HPI-mediated inhibition and allow for comparative analysis of inhibitor-bound enzymes, in addition to the pritelivir-bound H/P complex, we determined structures of H/P bound to IM-250 (2.9 Å) and amenamevir (2.8 Å) (Figures S4b and S5). All three inhibitors bind to a pocket that includes residues at the interface of the UL5 1A domain, 2A domain, and CTD, and the UL52 NTD and middle domains (Figures 3a, 3c, and 3e).

**Figure 3.**
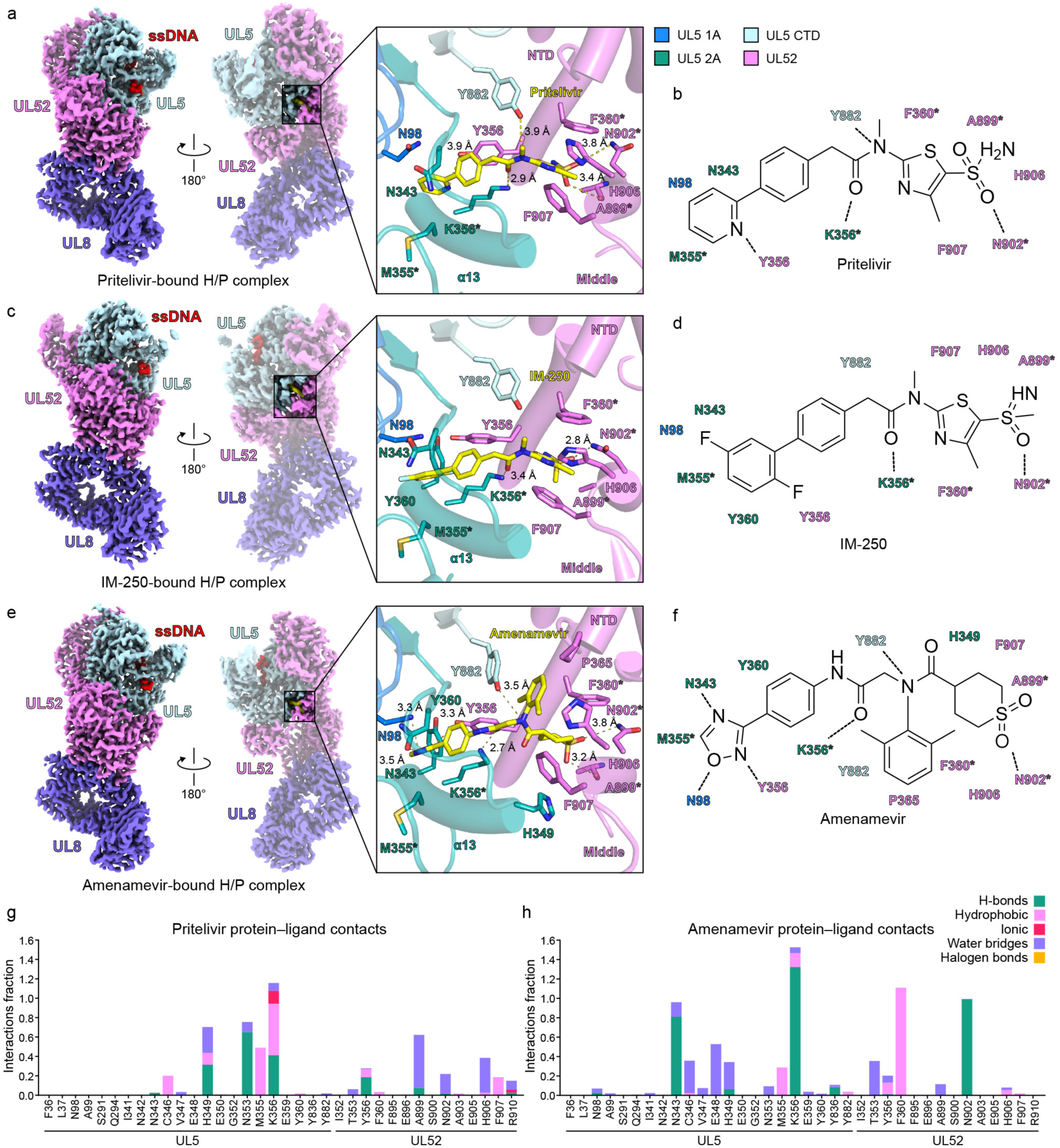
HSV H/P inhibitor binding modes. (a, c, and e) Cryo-EM maps (left) and ribbon diagrams (right) of HSV H/P complexes bound to inhibitor pritelivir (a), IM-250 (c), or amenamevir (e). UL5 domains are colored as indicated. Inhibitor-interacting residues are shown as sticks. The inhibitors are shown as yellow sticks. (b, d, and f) Schematic representations of the interactions between the HSV H/P complexes and inhibitors. Dashed lines indicate polar contacts. (g and h) Interaction fraction analysis between the H/P complex and pritelivir (g) or amenamevir (h) based on molecular dynamics simulation. H-bonds, hydrogen bonds; hydrophobic, non-polar contacts; ionic, electrostatic interactions; water bridges, mediated by water molecules; halogen bonds, involving halogen atoms in the ligand. In panels a–f, residues that are sites of drug-resistance-associated substitutions are labeled with asterisks. See Table S5 for additional information.

Pritelivir is in an extended conformation when it binds the H/P complex (Figures 3a and S1f). The 2-pyridyl moiety inserts into a pocket between the UL5 1A and 2A domains, while the aminosulfonyl group occupies a depression formed by α-helices from the UL52 NTD and middle domains. Pritelivir makes hydrophobic contacts with UL5 residues N98, M355, and Y882 and with UL52 residues Y356, F360, A899, H906, and F907. Pritelivir makes polar contacts with UL5 residues N343, K356, and Y882 and with UL52 residue N902 (Figure 3b).

In IM-250, the original 2-pyridyl ring in pritelivir was replaced with a 2,5-difluorophenyl group, and the sulfonamide group in pritelivir was replaced with a methylsulfoximine moiety (Figures 3c and 3d).^33^ The fluorinated phenyl moiety inserts into the same UL5 1A/2A cleft as the pritelivir pyridine ring, and the methylsulfoximine moiety fits into the UL52 depression formed by the α-helices of the UL52 NTD and middle domains, like the sulfonamide in pritelivir. The overall similar binding mode of both inhibitors is expected, and the agents have similar potency in antiviral assays.^33^

Amenamevir has a three-lobed architecture (Figures 3e, 3f, and S5e). Its oxadiazole group occupies the same UL5 pocket as the pyridine ring of pritelivir, while its thiane 1,1-dioxide ring fits into the same UL52 depression as the aminosulfonyl group of pritelivir. A third moiety, a 2,6-xylyl group, fits into a hydrophobic cleft formed by the side chains of UL5 CTD residue Y882 and UL52 NTD residue F360 (Figure 3f).

To further evaluate the importance of inhibitor–protein interactions identified in the cryo-EM structures, we conducted all-atom molecular dynamics (MD) simulations of the HSV H/P complex bound to pritelivir and amenamevir. In both simulations, UL5 residue K356 had the highest interaction fraction for both antivirals, involving hydrophobic and hydrogen-bonding interactions (Figures 3g and 3h). UL5 K356 thus likely plays a major role in stabilizing drug binding. UL5 N343 and Y882, and UL52 Y356, F360, and N902 also made important contributions to engaging both drugs.

### Determinants of HPI selectivity

Seven UL5 residues contact HPIs (Figure S6a). UL5 contact residues are highly conserved across herpesviruses; for example, UL5 residues N343 and K356, which the MD simulations suggest are important for drug binding, are uniformly conserved in α-, β- and γ-herpesviruses. Seven UL52 residues contact HPIs (Figure S6b). UL52 Y356, which is the UL52 residue with the highest interaction fraction with HPIs (Figures 3g and 3h), is also conserved among herpesviruses. However, UL52 contact residues F360, P365, A899, N902, H906, and F907 show some conservation among α-herpesviruses but are mostly not conserved in β-, and γ-herpesviruses. This divergence in the UL52 binding region likely accounts for the lack of activity of pritelivir and amenamevir against β-herpesviruses.^45^

Interestingly, amenamevir is active against HSV and VZV.^36,37,46,47^ The inhibitor’s cross-reactivity between HSV and VZV can be attributed to its unique three-lobed architecture, which includes the 2,6-xylyl group that provides contacts at the UL5–UL52 interface that pritelivir does not. The 2,6-xylyl group interacts with the side chains of UL5 Y882 and UL52 F360, which are well conserved within α-herpesviruses. Interactions with these conserved residues may compensate for potentially weakened interactions between amenamevir and other, less conserved regions in UL52.

### Determinants of HPI resistance

Among the previously identified HPI-resistance mutations, several residues directly contribute to drug binding (Table S5)^33,47–55^. These include M355I/T and K356N/T/Q in the UL5 2A domain, F360C/V in the UL52 NTD, and A899T and N902T in the UL52 middle domain. Mutations at these sites likely compromise inhibitor binding by altering direct interactions with HPIs (Figures 3g and 3h). Additionally, UL5 2A domain substitutions N342K and G352C/V/R, although they do not directly affect HPI contact residues, are adjacent to the HPI binding site and may cause steric hindrance or reorganization of pocket residues, thereby impairing inhibitor binding (Figure S6c). Furthermore, two amenamevir-resistance mutations, UL52 NTD substitutions S364G and R367H, are located close to the 2,6-xylyl group of amenamevir but are much more distant (7.5 Å) from the pritelivir and IM-250 binding site. This observation is consistent with a more prominent role for the UL52 NTD in amenamevir binding than in interactions with the other two HPIs.

### Structural basis for H/P complex inhibition

In SF1 helicases, ATP-dependent conformational cycling between two RecA-like domains, 1A and 2A, drives translocation along ssDNA.^12,39^ Structures of SF1 family members have shown that nucleotide binding induces the RecA domains to transition from an open to a closed conformation, facilitating directional movement along the DNA backbone^39^. Consistent with this general mechanism, the HSV UL5 RecA domains are expected to transition from an open to a closed conformation during translocation. To understand how HPIs inhibit DNA unwinding, we compared the HPI-bound cryo-EM structures to an AF3-predicted^40^ model of the ATP-bound, closed conformation (Figure S4a).

The inhibitors bind at the interface between UL5 domains 1A and 2A (Figure 4a). This binding mode blocks closure of the 2A domain towards the 1A domain, a conformational change necessary for DNA translocation. Structural alignment with the AF3-predicted ATP-bound H/P complex reveals displacement of two helices within the 2A domain—α13 and α29—that normally approach the 1A domain upon ATP binding. In the pritelivir-bound structure, α13 and α29 are shifted away from their ATP-bound positions by 2.8 Å and 3.6 Å, respectively (Figure 4a). Similar conformational differences are observed in the IM-25 and amenamevir-bound complexes (Figures 4b and 4c), indicating that all three inhibitors likely lock the RecA domains in an inactive, open configuration.

**Figure 4.**
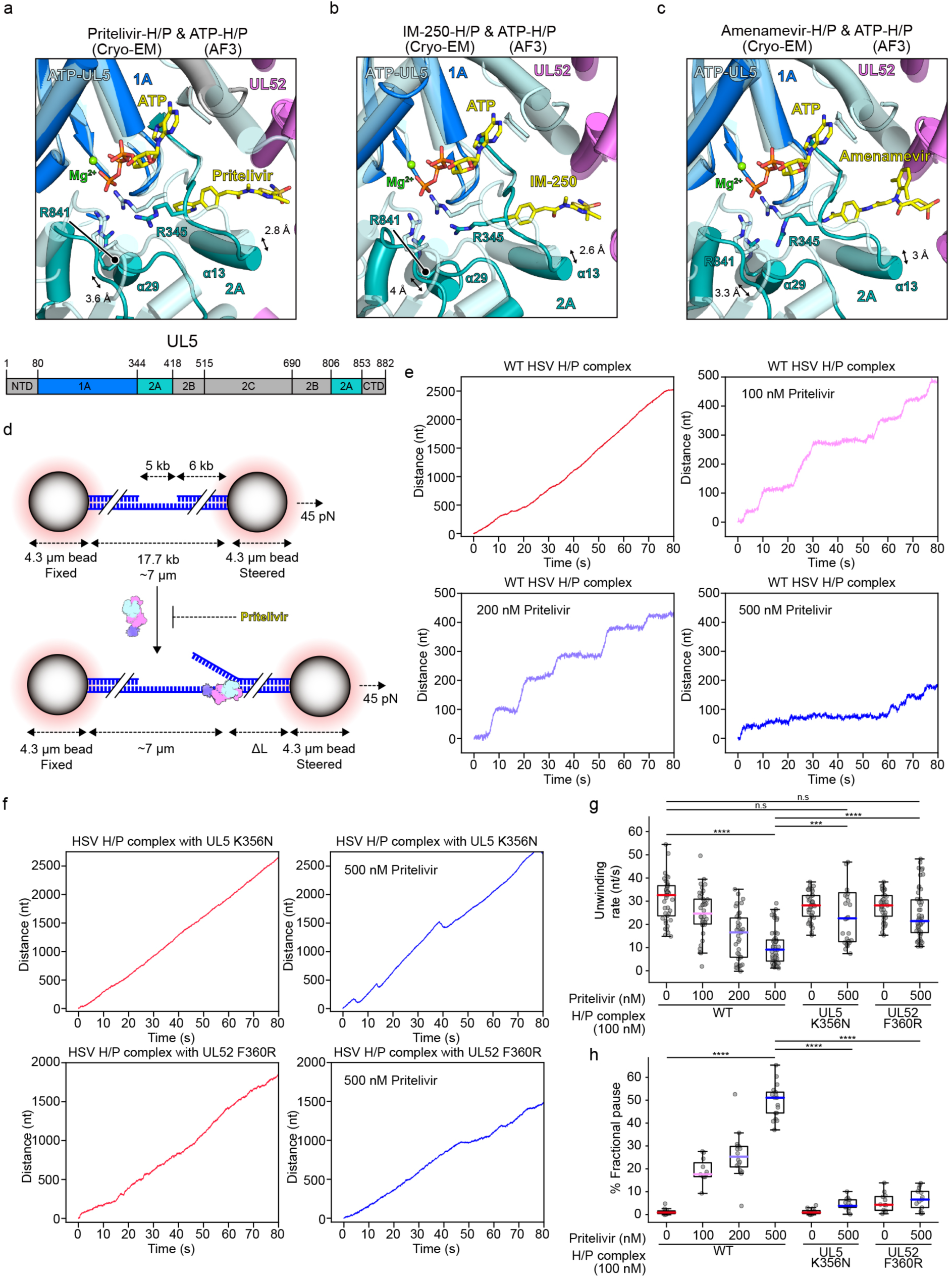
HPIs lock the H/P complex in an inactive conformation and cause it to pause on DNA. (a–c) Superposition of AF3^40^-predicted ssDNA- and ATP-bound HSV H/P complex with the cryo-EM structures of ssDNA- and pritelivir-bound complex (a), ssDNA- and IM-250-bound complex (b), and ssDNA- and amenamevir-bound complex (c). (d) Schematic diagram of the optical tweezer experiments. Biotinylated hybrid dsDNA/ssDNA molecules were suspended by two streptavidin-coated beads manipulated by optical traps. DNA length is expected to increase under a constant force (45 pN) when dsDNA is converted to ssDNA by helicase activity. (e) Representative traces of unwinding activity (from left to right) in the presence of 100 nM wild-type (WT) HSV H/P complex alone, or in the presence of pritelivir at the indicated concentrations. (f) Representative traces of unwinding activity (from left to right) of 100 nM HSV H/P complexes containing the UL5 K356N or UL52 F360R mutant subunits alone or in the presence of 500 nM pritelivir. (g) Comparison of DNA unwinding rates under various conditions with WT or mutant H/P complexes. Colored lines represent median values for different conditions. Statistical comparisons between experimental conditions were performed using Welch’s t-test. \*\*\*\**P* < 0.0001; not significant (n.s.); WT H/P complex with 500 nM pritelivir vs. UL5 K356N with 500 nM pritelivir, \*\*\**P* = 0.00246. (h) Comparison of the fractional pause state of conditions described in panel G. Statistical comparisons between experimental conditions were performed using Welch’s t-test (\*\*\*\**P* < 0.0001).

Examination of HPI-bound structures also reveals that inhibitor binding may impair positioning of the two arginine residues in the UL5 2A domain that coordinate the γ-phosphate of ATP (R345 and R841). R345 and R841 would be displaced due to the widened 1A–2A interface, suggesting that drug binding may impair ATP hydrolysis (Figures 4a–c).

The inhibitor-bound structures, therefore, suggest that HPIs likely allosterically inhibit helicase activity and lockUL5 in an open conformation by preventing ATP-induced closure of the RecA domains required for translocation on DNA.

### HPIs cause H/P complexes to pause on DNA

To test the hypothesis that HPIs lock H/P complexes in an inactive conformation, we next turned to single-molecule experiments performed with optical tweezers. To monitor H/P activity on individual DNA molecules in the presence of HPIs, we used a dual-trap optical tweezers assay with a 17.7 kb dsDNA/ssDNA hybrid substrate tethered between two streptavidin-coated polystyrene beads (Figure 4d).^56,57^ We conducted unwinding assays in constant-force mode, where a high-frequency feedback system maintained a force set point of 45 pN. In these assays, DNA unwinding results in increased bead-to-bead distance due to the conversion of dsDNA into ssDNA.^58^

In the presence of WT HSV H/P complex alone, we observed robust and continuous DNA unwinding, with no detectable pausing, with an average rate of 33 nt s⁻¹ (*n* = 41). Upon addition of pritelivir, unwinding activity was progressively reduced in a concentration-dependent manner, and concomitantly, the H/P complex spent more time in the paused state during unwinding (Figures 4e, 4g, 4h, and S7a).

At 100 nM pritelivir, the average unwinding rate decreased to 25 nt s⁻¹ (*n* = 38), with a corresponding pause fraction of 17.9% (*n* = 11). At 200 nM, the rate further declined to 16.8 nt s⁻¹ (*n* = 37), and the pause fraction increased to 25.6% (*n* = 15). At the highest concentration tested, 500 nM, unwinding activity was severely impaired, with an average rate of only 9.5 nt s⁻¹ (*n* = 51) and a pause fraction of 51.5% (*n* = 16) (Figures 4e, 4g, 4h, S7b, and S8). These results indicate that pritelivir slows the helicase translocation rate and promotes pausing in a dose-dependent manner, consistent with stabilization of an inactive conformation of the H/P complex.

To better understand the drug selectivity of HPIs, we introduced two point mutations at the inhibitor binding site: UL5 K356N and UL52 F360R. UL5 K356N is a well-characterized resistance mutation (Table S5)^50,55,59,60^ and K356 is the UL5 residue that makes the most HPI contacts as observed in MD simulations (Figures 3g and 3h). UL52 F360R replaces a phenylalanine that is conserved in α-herpesviruses with the naturally occurring arginine found in HCMV (Figure S6b). This substitution disrupts a key hydrophobic interaction between UL52 F360 and the inhibitor, and we hypothesized that it may, at least in part, explain why pritelivir is not active against HCMV.

Consistent with drug resistance, H/P complexes containing either the UL5 K356N or UL52 F360R mutant subunits had helicase activity in the presence of 500 nM pritelivir. The UL5 K356N mutant had an unwinding rate of 23 nt s⁻¹ (*n* = 32) with only a 4% pause fraction (*n* = 13), while the UL52 F360R mutant had an unwinding rate of 22 nt s⁻¹ (*n* = 50) and a 6.8% pause fraction (*n* = 12). When tested in the absence of pritelivir, both mutants had unwinding kinetics comparable to the WT H/P complexes (Figures 4f–h, S9, and S10), indicating that these mutations do not intrinsically impair helicase function. Thus, the UL52 F360R substitution that naturally occurs in HCMV UL70 may explain at least in part why this β-herpesvirus is not susceptible to pritelivir.

### Replication fork complex assembly

We purified the HSV polymerase holoenzyme (UL30, residues 43–1235, and UL42, residues 2–340) from insect and bacterial cells, respectively. Mixing the H/P complex with polymerase in the absence of DNA did not yield a substantial population of particles that were larger than H/P complex (Figure S11a and Table S1). However, co-incubating the H/P complex with polymerase, a DNA replication fork substrate, and pritelivir resulted in an assembly that was stable when analyzed by SEC. Sodium dodecyl-sulfate polyacrylamide gel electrophoresis (SDS–PAGE) analysis of the SEC peak sample revealed bands of the expected sizes for subunits of the H/P complex and polymerase holoenzyme (Figure S11b). Analysis of the SEC peak sample by MP revealed a large proportion of particles with a mass of 494 kDa, which is close to the expected mass of one copy each of the H/P complex, polymerase holoenzyme, and DNA substrate (Figure 5a and Table S1).

**Figure 5.**
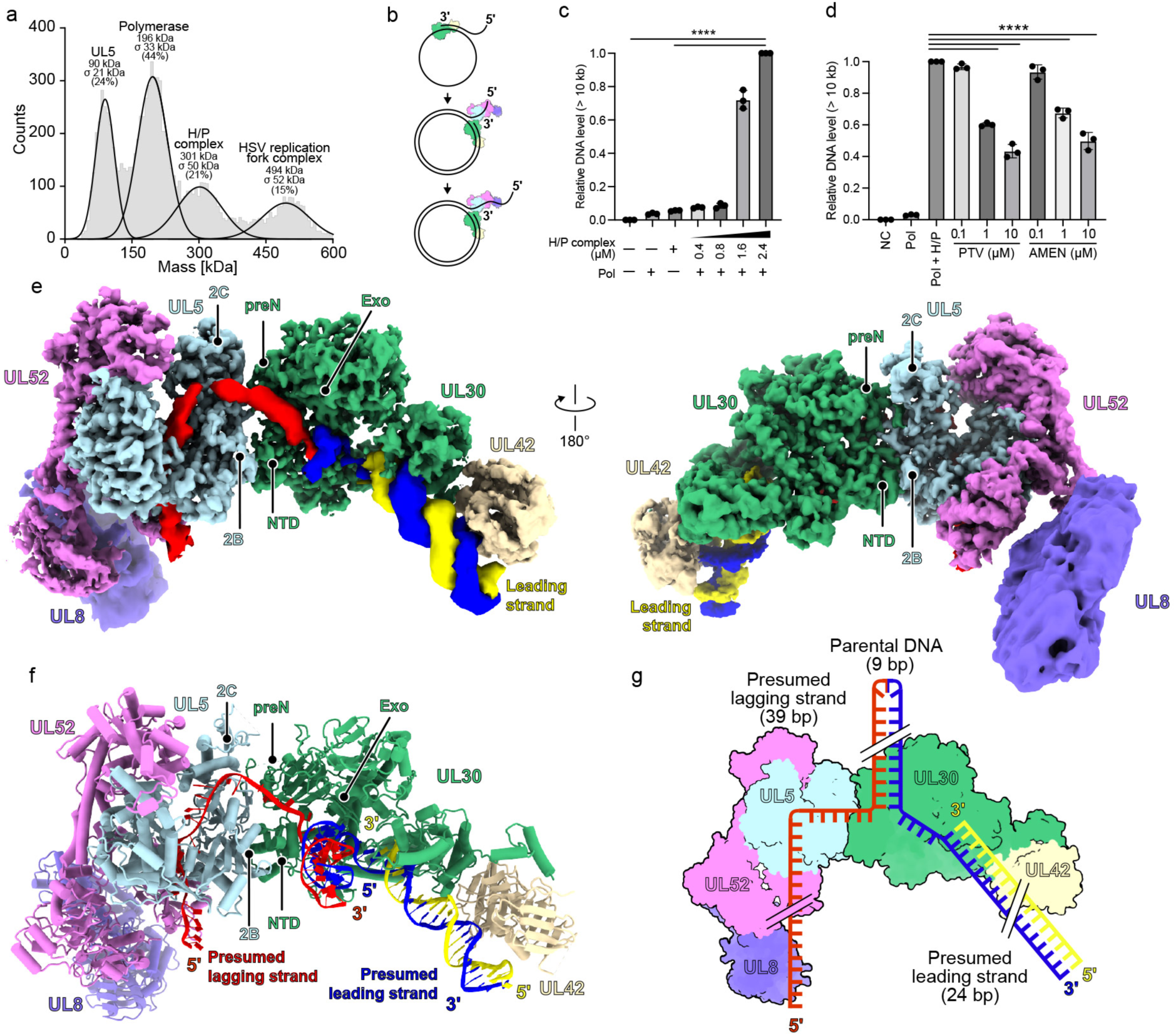
Structure of HSV replication fork complex bound to DNA. (a) Mass photometry results of SEC-purified HSV replication fork complex. Based on expected masses, the 494 kDa peak likely represents the HSV replication fork complex, the 301 kDa peak likely represents the H/P complex, the 196 kDa likely represents the polymerase holoenzyme, and the 90 kDa peak likely represents UL5. This experiment was performed twice, with representative data shown. Please see Table S1 for additional information. (b) Schematic diagram of the rolling-circle assay. The polymerase holoenzyme is shown in green and yellow. The H/P complex is shown in blue, pink, and purple. The assay is also performed in the presence of the HSV-1 single-stranded DNA-binding protein, ICP8 (not shown). (c) Quantification of the DNA products (> 10 kb DNA, see SFig. 11c) from the rolling circle assay performed with HSV polymerase with or without co-incubation with H/P complex. Mean ± standard deviation from three independent experiments. One-way ANOVA with Dunnett’s multiple comparison test (\*\*\*\**P* < 0.0001). (d) Quantification of the levels of DNA products (> 10 kb DNA, see SFig. 11d) for rolling-circle assay performed in the presence of pritelivir (PTV) or amenamevir (AMEN). NC (negative control): annealed M13mp18 and primer only; PC (positive control): 0.4 μM Pol + 2.4 μM H/P complex, no HPI; mean ± standard deviation from three independent experiments. One-way ANOVA with Dunnett’s multiple comparison test (\*\*\*\**P* < 0.0001). (e and f) Cryo-EM map (e) and ribbon diagram (f) of the HSV replication fork complex bound to fork DNA, colored as indicated. UL5 and UL30 domains involved in H/P and polymerase interactions are indicated. Two views are shown. (g) Schematic diagram of the fork DNA-bound HSV replication fork complex.

We performed a DNA replication assay using a substrate comprised of a circular M13 ssDNA template annealed to an oligonucleotide (Figure 5b and Table S2). Reactions were assembled with a constant concentration of the single-stranded DNA binding protein ICP8 (0.8 μM) and polymerase holoenzyme (UL30–UL42) (0.4 μM) and increasing concentrations of the H/P complex (0.4–2.4 μM). In the absence of H/P complex, only template-length products (∼7 kb) were observed, reflecting extension by the polymerase alone until it reached a region of dsDNA. Upon addition of the H/P complex, however, products exceeding 10 kb were generated in an H/P complex-concentration-dependent manner, indicating that helicase activity enabled DNA unwinding, promoting replication through dsDNA. Higher H/P concentrations progressively enhanced long-product synthesis, indicating that helicase activity promotes DNA unwinding and processive replication (Figures 5c and S11c).

To assess the effects of HPIs, we performed these DNA replication assays in the presence of increasing concentrations of pritelivir and amenamevir (Figures 5d and S11d). After adding these inhibitors, the amounts of long DNA products were markedly reduced in a dose-dependent manner, indicating efficient inhibition of the HSV replication fork complex. Importantly, the synthesis of the 7 kb product, which is primarily generated by DNA polymerase holoenzyme in the absence of helicase activity, remained unaffected (Figure S11d). These findings suggest that HPIs specifically inhibit the H/P complex in the HSV replication fork, as expected, without impairing the catalytic activity of the polymerase.

### Overall structure of the HSV DNA replication fork complex

We used single-particle cryo-EM to obtain a map of a replication fork complex comprising pritelivir-inhibited HSV H/P complex described above, polymerase holoenzyme, and fork DNA, at a global resolution of 3.8 Å. We used masked refinement to visualize and build various portions of the complex (Figures S11e and Table S3). Map quality was the poorest for UL8 and UL42, likely due to flexibility. To model these regions of the complex, we therefore used the high-resolution structure of the pritelivir-bound H/P complex and previously reported structures of the HSV polymerase holoenzyme.^17,24,26^

The HSV replication fork complex has an elongated architecture, with UL5 enveloping the ssDNA (presumed lagging strand) and UL5 interacting with UL30 (Figures 5e and 5f). This assembly is organized around a 39-nt ssDNA strand and a 24-bp presumed leading-strand duplex that includes the primer for the polymerase (Figure 5g). The DNA duplex is situated in the center of the complex, near a cleft formed by the UL30 Exo domain and NTD (Figure S12a), which is adjacent to the UL30 ssDNA binding groove.^26,27^ Although the flexibility of the parental DNA precluded high-resolution modeling, we observed clear density for the DNA duplex in the low-contour map near a pocket formed by the NTD and Exo domain of UL30 (Figure S12a). This positioning implies that the UL30 NTD may contribute to coordinating polymerase and helicase activities by stabilizing the parental DNA at the replication fork.

### DNA interactions in the replication fork complex

Assembly of the HSV replication fork complex generates a positively charged groove that guides the ssDNA strand through the interface of the H/P complex and polymerase (Figure S12b). These DNA-interacting residues form a continuous path involving extensive polar, electrostatic, and hydrophobic interactions (Figure S12c). The ssDNA strand enters this channel from the fork junction and is initially clamped between the UL30 NTD and the UL5 2B and 2C domains (Figures 6a and 6b). UL30 residues H158, Y160, R163, and R496 make hydrogen bonding and stacking interactions with the DNA, directing the ssDNA toward the helicase surface. UL52 residues R547, R713, and F737 interact with the phosphate backbone and bases of the ssDNA within the positively charged groove (Figure 6b).

**Figure 6.**
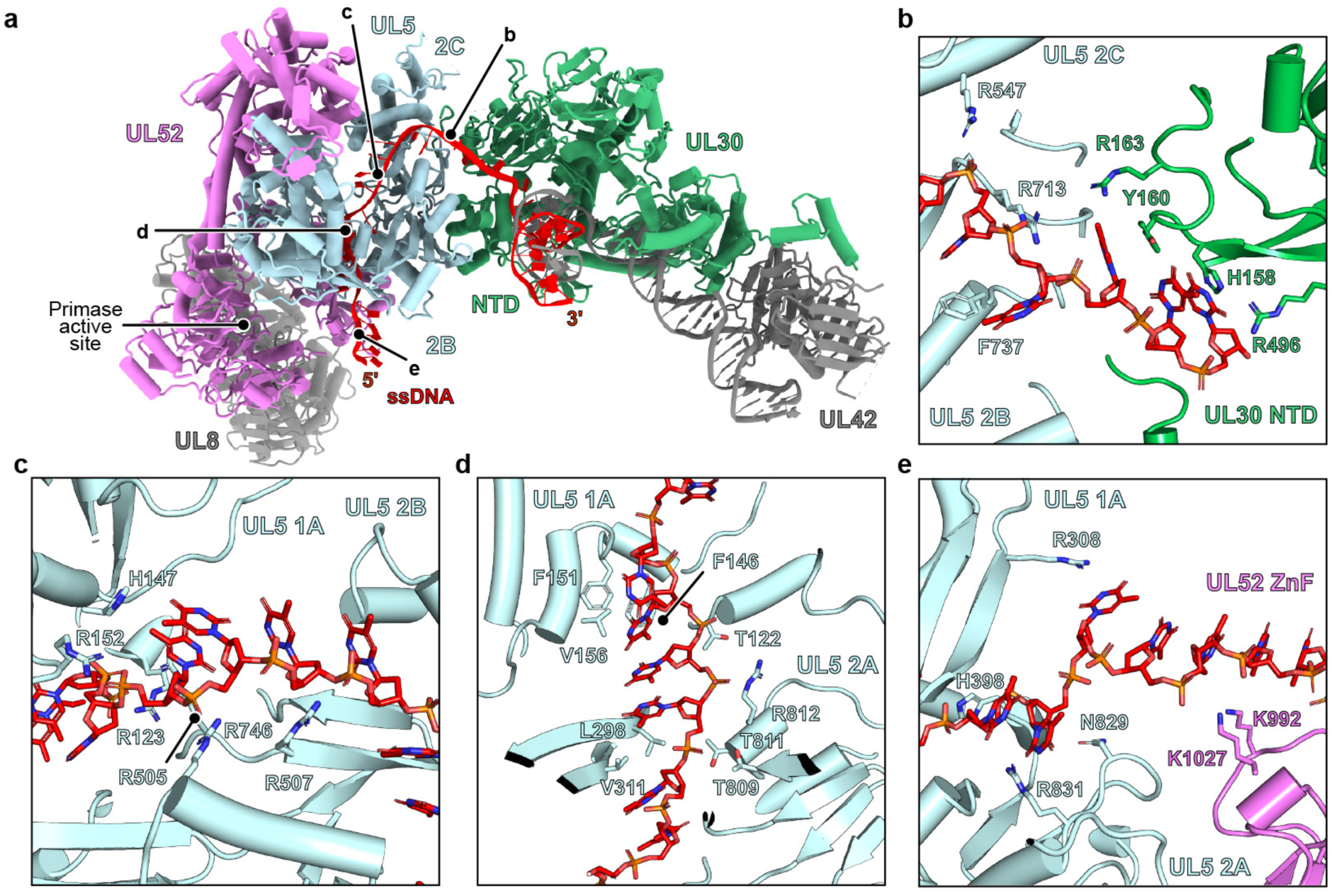
Replication fork complex interactions with DNA. (a) Ribbon diagram of the HSV replication fork complex bound to DNA. The general locations of the views shown in panels B–E and the UL52 primase active site are indicated. (b) UL30 NTD and UL5 2B and 2C domain interactions with DNA. (c) UL5 1A and 2B domain interactions with DNA. (d) UL5 1A and 2A domain interactions with DNA. (e) UL5 1A and 2A domains and UL52 ZnF domain interactions with DNA.

The ssDNA then traverses a path across the UL5 helicase. A network of basic residues lines the outer face of the UL5 1A and 2B domains, making interactions with the DNA backbone (Figure 6c). Additional polar and hydrophobic contacts bind the ssDNA in a cleft formed by the helicase 1A and 2A domains (Figure 6d). These residues span the DNA path along the 1A and 2A RecA-like domains, forming a continuous groove that positions the DNA for 5′–3′ translocation. There is a final set of electrostatic and hydrogen-bonding interactions as the ssDNA exits UL5 at the distal end of the groove, with adjacent residues from the UL52 ZnF, K992 and K1027, interacting with the ssDNA (Figure 6e). An unexpected feature is that, despite the ssDNA segment in the DNA substrate we designed being long enough to reach the UL52 primase active site, we did not observe it reach this site.

### FYNPYL motif mediates replication fork complex assembly

The preN domain of UL30, comprising residues 1–140, is a herpesvirus-specific structural feature.^17^ This domain contains a highly conserved FYNPYL motif and has been suggested to be involved in protein–protein interactions important for viral DNA replication, although mutating it does not impair UL30 5′–3′ polymerase activity (Table S4).^18,19^ In the replication fork complex, the preN FYNPYL motif extends towards UL5 and interacts with the UL5 2B and 2C domains (Figures 7a and 7b). Although the resolution in this portion of the structure was low (in the 4.1 Å focused map: see Figure S13a), interactions between the FYNPYL motif and UL5 could also be predicted by AF3.^40^ Specific contacts include those made by F44, Y48 and L49, in the UL30 FYNPYL motif with UL5 residues F550, Y648, and Y696, which are conserved in herpesviruses (Figures 7d and 7e). Additionally, the side chain of UL5 N699 interacts with the side chain of UL30 Y45 and the backbone carbonyl of UL30 N46 (Figure 7b).

**Figure 7.**
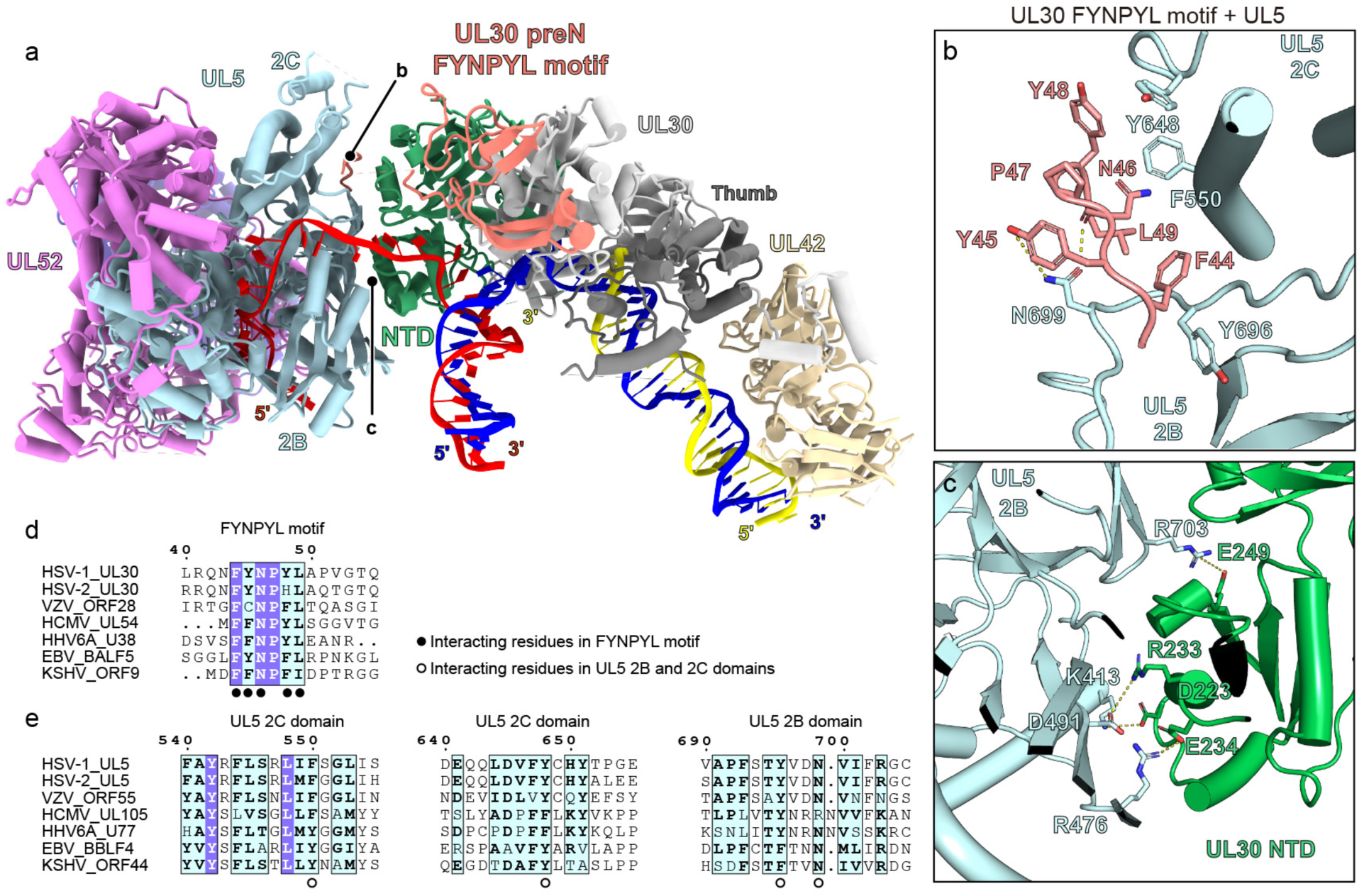
Herpesvirus-specific FYNPYL motif participates in replication fork complex assembly. (a) Ribbon diagram of the fork DNA-bound HSV replication fork complex. The UL30 preN domain, shown in salmon, contains the conserved FYNPYL motif that interacts with the UL5 2C and 2B domains. The general location of the views shown in panels B and C are indicated. (b) Interactions between the HSV UL30 FYNPYL motif and UL5 2B and 2C domains. (c) Interactions between the HSV polymerase UL30 NTD and helicase UL5 2B domain. (d–e) Sequence alignment of FYNPYL motif in the UL30 (polymerase catalytic subunit) preN domain (d), or UL5 2B and 2D domain regions that interact with the UL30 NTD (e). Conserved residues are indicated. HSV-1, herpes simplex virus 1; HSV-2, herpes simplex virus 2; VZV, varicella-zoster virus; HCMV, human cytomegalovirus; EBV, Epstein–Barr virus; HHV-6A, human herpesvirus 6A; KSHV, Kaposi’s sarcoma-associated herpesvirus.

Sequence alignments across herpesvirus polymerase and helicase confirm the conservation of these residues (Figures 7d and 7e). Comparison of AF3^40^-predicted models of α-, β-, and γ-herpesvirus H/P complexes with the residues comprising the motif from each virus suggests evolutionary preservation of this interaction (Figures S13b–m).

In addition to the UL30 preN domain and the UL5 2B/2C domains, we identified extensive polar interactions between the UL30 NTD and the UL5 2B domain (Figure 7c). These interactions likely further stabilize the UL30–UL5 interface. However, sequence alignments suggest that this interface is not conserved across herpesvirus species and may form through different sets of contacts in the other viruses (Figure S14).

## Discussion

Coordinated actions of the H/P complex and the DNA polymerase holoenzyme drive herpesvirus DNA replication. While the enzymatic functions of these components are well established, the structural basis underlying their coordination and regulation has remained unclear. In this study, in addition to defining how the H/P complex is selectively targeted by HPIs, we clarify how the HSV replication fork assembles on DNA.

The cryo-EM structures reveal that HPIs lock UL5 in an open, catalytically inactive conformation. Each inhibitor binds at a cleft formed by the UL5 1A and 2A domains, preventing conformational closure required for ATP hydrolysis and DNA translocation. This allosteric inhibition mechanism, which targets a conformational state rather than the helicase or primase active sites, offers a distinct approach from nucleoside analogs targeting the polymerase UL30.

Despite their conserved mode of action, current HPIs show efficacy almost exclusively against α-herpesviruses. Our structural data provide a likely explanation: while the UL5-interacting residues that form the core of the inhibitor binding site are well conserved, the adjacent residues contributed by UL52—which shape the periphery of the HPI binding site—are more variable across herpesvirus subfamilies. This divergence likely underlies the lack of activity of HPIs against the β-herpesvirus HCMV. Given the expected structural conservation of the H/P complex across herpesviruses, HPIs rationally designed to target the same binding site in the HCMV H/P complex could, in principle, be developed to achieve the same mode of action. The structures thus suggest a path toward developing HPIs that are selective against HCMV. They suggest that developing broader spectrum agents targeting the same site may be challenging, given sequence divergence in the UL52 region.

The cryo-EM structure HSV replication fork complex reveals a herpesvirus-specific interface linking the polymerase UL30 and helicase UL5, mediated by the conserved FYNPYL motif within the UL30 preN domain and the 2B/2C domains of UL5. This interaction directly links the polymerase and helicase modules and likely facilitates the coordination of DNA unwinding with strand synthesis. Notably, both the polymerase FYNPYL motif and the UL5 2C domain are unique to herpesviruses, suggesting that this interface serves as a virus-specific module that promotes efficient replication. Interestingly, recombinant HSV-1 in which the FYNPYL motif is replaced by six alanine residues (*pol*A_6_) when studied a mouse model of corneal infection causes a modest defect at the site of inoculation but results in a severe defect in ganglionic replication.^19^ It was thus hypothesized that the motif interacts with a host factor that may be differentially expressed in different cell types or tissues. However, our findings suggest that part of the replication defect likely arises from the ability of the motif to interact with UL5. Indeed, in addition to the FYNPYL–-UL5 interaction interface, we also observed additional contacts between the UL30 NTD and UL5 2B domain that would allow the polymerase and helicase to maintain some interactions with each other even if the UL30 preN FYNPYL motif–UL5 interaction is disrupted.

One unexpected feature of the cryo-EM structure of the DNA-bound replication fork complex is that the ssDNA does not reach the UL52 primase AEP site, despite the fact that we included an oligonucleotide that should have been long enough to do so (e.g., the 5′ overhang from that segment, not visualized due to flexibility, is 18 nucleotides long) (Figure 6a). Future experiments with modified DNA substrates, including multiple primase DNA sequence recognition sites^61^, may be required to determine whether this would allow visualization of ssDNA in the primase active site.

In summary, this study elucidates the mechanism by which HPIs block helicase function by locking the H/P complex in an inactive conformation, causing the helicase to pause on DNA. These insights advance our understanding of viral replisome architecture and provide a structural framework for developing antivirals targeting essential functions of the herpesvirus replication machinery.

## Acknowledgements

Cryo-EM data were collected at the Harvard Cryo-EM Center for Structural Biology at Harvard Medical School. We would like to thank Richard Walsh, Megan Mayer, Conny Leistner, Remya Nair, and Shaun Rawson at the Harvard Cryo-EM Center. This work was supported by NIH awards R21 AI141940 to D.M.C. and J.A, R01 AI019838 to D.M.C. and James Hogle, R01 AI021747 to S.K.W., and R01 CA272436 to J.J.L. This work was also supported by a grant from the Broad Institute Center for Integrated Solutions for Infectious Diseases (CISID) to J.A. J.A. is a recipient of a Burroughs Wellcome Fund Investigator in the Pathogenesis of Infectious Disease Award and is a Howard Hughes Medical Institute Investigator. We thank Dr. Bridget Gollan for help with editing figure illustrations.

## Contributions

Z.Y. designed and cloned the HSV H/P complex constructs and designed DNA oligos, produced recombinant proteins, carried out rolling-circle assays, performed optical tweezer and mass photometry experiments, collected and processed cryo-EM data, built and refined atomic models of HPI-bound HSV H/P complexes and HSV replication fork complex, wrote the initial draft of the paper, and generated figures. P.S. performed optical tweezer experiments and analyzed and interpreted data. C.L. performed the MD simulation. P.Y. produced HSV UL30. D.M.C. and S.K.W. participated in study design. M.S. helped design and supervised the MD simulation. J.J.L. supervised the optical tweezer experiments and acquired funding. J.A. supervised the study, acquired funding, and revised and edited the manuscript. All authors participated in reviewing and editing the manuscript.

## Methods

### Protein expression and purification

Full-length open reading frames (ORFs) of three subunits, UL5 (residues 1–882; GenBank YP_009137079.1), UL52 (residues 1–1058; GenBank YP_009137128.1), UL8 (residues 1–750; GenBank YP_009137082.1) of the HSV H/P complex were subcloned into a modified pCAG vector with or without a maltose binding protein (MBP) tag at the N terminus. We used PCR-based mutagenesis, with the plasmids noted above as starting points, to generate the UL5 K356N and UL52 F360R mutants.

All subunits were co-transfected into suspension Expi293F cells using PEI (25,000 MW, Polysciences) when the cells reached a density of 2 × 10^6^ cells ml^-1^, in a 6:3:1 UL5/UL52/UL8 ratio. The purified complexes were subjected to SDS–PAGE analysis (Figure S1a).

After culturing at 37 °C for 72 h, cells were harvested by centrifugation at 4000 rpm for 20 min. Cells were lysed using lysis buffer (50 mM HEPES-NaOH, pH 7.5, 300 mM NaCl, 0.5% (v/v) Triton X-100, 5 mM MgCl_2_, 0.5 mM EDTA, 1 mM DTT, and protease inhibitor [cOmplete™, Mini, EDTA-free protease inhibitor cocktail, Millipore Sigma, Cat# 11836170001]). Cell debris was removed through centrifugation 50,000 x g for 2 h with Ti50.2 rotor. The supernatant was incubated with amylose resin (NEB Cat# E8021S) at 4 °C for 1 h and washed with wash buffer (25 mM HEPES-NaOH, pH 7.5, 300 mM NaCl, 5 mM MgCl_2_, 0.5 mM EDTA, and 1 mM DTT).

Immunoprecipitated HSV H/P complex was subjected to on-column digestion overnight with HRV-3C protease (TaKaRa Cat# 7360). The eluate fractions within the elution buffer (25 mM HEPES-NaOH, pH 7.5, 150 mM NaCl, 2 mM MgCl_2_ and 1 mM DTT) were concentrated at 1.1 mg ml^-1^ for further purification using a Superdex 200 Increase (Cytiva Cat# 28990944) SEC column. Fractions containing HSV H/P complex were pooled, concentrated to ∼0.3 μg ml^-1^, and flash-frozen in liquid nitrogen or used for subsequent analysis. At each purification step, we monitored proteins for purity and stoichiometry by SDS–PAGE analysis (Figure S1a).

To investigate the function of key HPI-binding residues within the HSV H/P complex, we purified two mutant complexes for optical tweezer experiments. In the first mutant complex, a point mutation (K356N) was introduced into the UL5 subunit, while the UL8 and UL52 subunits remained WT. In the second mutant complex, a point mutation (F360R) was introduced into the UL52 subunit, with WT UL5 and UL8. Both mutant complexes were co-expressed in Expi293F cells and purified following the same protocol as for the WT complex.

Wild-type KOS strain HSV UL30 (GenBank: AFE62858.1) containing a 42-amino acid N-terminal deletion (HSV PolΔN42) and an N-terminal polyhistidine tag was expressed in insect cells (*Spodoptera frugiperda* Sf9 cells, ThermoFisher Scientific Cat# 11496015) using recombinant baculovirus and harvested 60–72 h post infection, and centrifuged at 2,800 x g for 30 min. We washed pellets twice in Dulbecco’s phosphate-buffered saline (DPBS) (Thermo Fisher Scientific Cat# 14190144) containing 10% (v/v) glycerol and froze pellets at -80 °C. After thawing prior to use, we resuspended cell pellets in buffer A (25 mM HEPES pH 7.5, 500 mM NaCl, 5 mM imidazole, 10% (w/v) sucrose) supplemented with protease inhibitor tablet (Millipore-Sigma Cat# 11836170001) and lysed them by sonication for 15 min (Branson Ultrasonics Sonifier Model S-450). We carried out all purification steps at 4 °C. Lysates were centrifuged at 30,000 x g for 1 h and filtered through a 0.45 μm membrane (MCE Membrane Filter) (Millipore Sigma Cat# HAWP14250). We then applied clarified supernatants onto a Hitrap™ Talon™ column (Cytiva Cat# 45-002-385) that was preequilibrated with buffer A. We washed the column with 20 column volumes of buffer A and eluted the protein using a gradient of 5 to 150 mM imidazole distributed over 20 column volumes. We collected fractions and analyzed these for purity by SDS-PAGE. Pol-containing fractions were then diluted 10-fold with H-buffer A (25mM HEPES pH 7.5, 1mM DTT, and 10% sucrose (w/v)) and loaded onto a HiTrap™ Heparin HP affinity column (Cytiva Cat# 45-000-057) preequilibrated with H-buffer B (H-buffer A supplemented with 0.1 M NaCl). We washed the column with 20 column volumes of H-buffer B and eluted with a gradient of 0.1 M to 1 M NaCl over 10 column volumes. We analyzed fractions using SDS-PAGE and pooled and concentrated fractions containing HSV UL30. We dialyzed these fractions using a Slide-A-Lyzer™ MINI Dialysis Device (20K MWCO) (Thermo Scientific Cat# 88405) into storage buffer (25 mM HEPES pH 7.5, 150 mM NaCl, 2 mM tris (2-carboxyethyl) phosphine (TCEP) (Oakwood Chemical Cat# M02624) and 20% (v/v) glycerol).

We expressed a UL42 (GenBank: AFE62870) construct containing an N-terminal MBP tag followed by a PreScission protease cleavage site and residues 1–339 of UL42 (MBP-UL42ΔC340) in *Escherichia coli* BL21 (DE3) pLysS cells (Novagen Cat# 69451) and purified the protein using amylose resin (NEB Cat# E8021S)^24,62^. Briefly, we grew cells that were transformed with MBP-UL42ΔC340 in the pMalc2 vector in Luria-Bertani (LB) broth with 2% (w/v) glucose in the presence of ampicillin at 100 µg ml^-1^ and chloramphenicol at 25 µg ml^-1^ for selection. Cells were induced with 0.3 mM isopropyl β-D-1-thiogalactopyranoside (IPTG) at OD_600_ ∼0.6–0.8 and allowed to grow overnight at 16 °C. We harvested cells by spinning down at 4000 x g for 30 min and resuspended them in lysis buffer (M-buffer A: 25 mM HEPES pH 7.5, 500 mM NaCl, 1 mM EDTA, 1mM DTT, and protease inhibitor (cOmplete™, Mini, EDTA-free protease inhibitor cocktail, Millipore Sigma, Cat# 11836170001)). We lysed cells by sonication for 15 min. We then centrifuged lysates for 90 min at 18,000 x g at 4 °C and applied supernatants to amylose resin (NEB Cat# E8021S). We washed the column with M-buffer A for 10–15 column volumes (CVs), eluted protein with M-buffer B (M-buffer A + 10 mM maltose) and analyzed fractions by SDS–PAGE. We pooled fractions and diluted these with H-buffer A to reduce the effective NaCl concentration to 150 mM. We cleaved the MBP-tag by adding polyhistidine-tagged HRV-3C protease (TaKaRa Cat# 7360) at 4 °C overnight and passed samples over a HiTrap TALON^®^ crude column (Cytiva Cat # 28953809) to remove the protease. We applied flowthrough from this step onto a HiTrap Heparin HP column (Cytiva Cat# 17040701) preequilibrated with H-buffer B. The column was washed with 20 CVs of H-buffer B and eluted with a gradient of 0.1M to 1M NaCl over 10 CVs. Fractions were pooled and stored in storage buffer.

Full-length HSV-1 ICP8 (GenBank: NC_024306.1) was subcloned into a modified pCAG vector with a maltose binding protein (MBP) tag at the N terminus. The plasmid was transfected into suspension Expi293F cells using PEI (25,000 MW, Polysciences) when the cells reached a density of 2 × 10^6^ cells ml^-1^. After culturing at 37 °C for 72 h, cells were harvested by centrifugation at 4000 rpm for 20 min. Cells were lysed using lysis buffer (50 mM HEPES-NaOH, pH 7.5, 500 mM NaCl, 0.5% (v/v) Triton X-100, 5 mM MgCl_2_, 0.5 mM EDTA, 1 mM DTT, and protease inhibitor [cOmplete™, Mini, EDTA-free protease inhibitor cocktail, Millipore Sigma, Cat# 11836170001]). Cell debris was removed through centrifugation 50,000 × g for 2 h with Ti50.2 rotor. The supernatant was incubated with amylose resin (NEB Cat# E8021S) at 4 °C for 1 h and washed with wash buffer (25 mM HEPES-NaOH, pH 7.5, 300 mM NaCl, 5 mM MgCl_2_, 0.5 mM EDTA, and 1 mM DTT). Resin-associated ICP8 was subjected to on-column digestion overnight with HRV-3C protease (TaKaRa Cat# 7360). The eluate fractions within the elution buffer (25 mM HEPES-NaOH, pH 7.5, 150 mM NaCl, 2 mM MgCl_2_ and 1 mM DTT) were concentrated for further purification using a Superdex 200 Increase (Cytiva Cat# 28990944) SEC column. Fractions containing ICP8 were pooled, and flash-frozen in liquid nitrogen or used for subsequent analysis.

### Cryo-EM sample preparation for HPI-bound helicase-primase

For HPI-bound H/P complex, the DNA template (5′-TCC CGC CCG TTT TTT TTT TTT TTT TTT TTT TTT TTT TTT TCG GAG TCG TTT CGA CTC CGA TTT TTA CAC GCT ATG TCG TCA AGT TGT ACC-3′) and primer (5′-GGT ACA ACT TGA CGA CAT AGC GTG-3′) were annealed by first incubating at 95 °C for 10 minutes and then slowly cooling samples to 4 °C (Table S2). The H/P complexes, DNA hybrid, and HPIs pritelivir (Sigma-Aldrich Cat# SML3422), IM-250 (Probe Chem Cat# PC-72690), or amenamevir (MedChemExpress Cat# HY-14809) were mixed at a 1:1.2:1.5 molar ratio in a buffer containing 25 mM HEPES-NaOH, pH 7.5, 150 mM NaCl, 2 mM MgCl_2_, and 1 mM DTT for 1 h for subsequent structural analysis, respectively.

All samples for cryo-EM were vitrified with a Vitrobot Mark IV (Thermo Fisher Scientific), with samples maintained at 100% humidity at room temperature. We applied 4 µl of sample to Quantifoil Au 1.2/1.3 300 mesh (EMS Q450CR1.3) grids that were previously plasma treated in a Pelco easiGlow^TM^ discharge cleaning system at 0.39 mBar, 15 mA, for 30 s, and used blot times of 3 s.

### Cryo-EM sample preparation for HPI-bound replication fork complexes

For HSV replication fork complex assembly, the DNA template (5′-TCC CGC CCG TTT TTT TTT TTT TTT TTT TTT TTT TTT TTT TCG GAG TCG TTT CGA CTC CGA TTT TTA CAC GCT ATG TCG TCA AGT TGT ACC-3′) and primer (5′-GGT ACA ACT TGA CGA CAT AGC GTG- 3′) were annealed by first incubating at 95 °C for 10 minutes and then slowly cooling to 4 °C in a (Table S2). The H/P complex, DNA-hybrid and pritelivir were mixed at 1:1.2:1.5 molar ratio in a buffer containing 25 mM HEPES-NaOH, pH 7.5, 150 mM NaCl, 2 mM MgCl_2_, and 1 mM DTT for 1 h. Following this, UL30, UL42, and dTTP (Invitrogen Cat# 10219012; at a molarity equal to H/P complex) were added and the sample was incubated at 4 °C overnight. The sample was passed on a Superdex 200 Increase (Cytiva Cat #28990944) SEC column. Fractions containing HSV replication fork complex were pooled and concentrated to ∼0.3 μg μl^-1^. For each purification step, proteins were monitored for purity and stoichiometry using SDS–PAGE analysis.

The sample of HSV replication fork complex was vitrified with a Vitrobot Mark IV (Thermo Fisher Scientific), with samples maintained at 100% humidity at room temperature. We applied 4 μl of sample to Quantifoil Au 1.2/1.3 300 mesh (EMS Q450CR1.3) grids that were previously plasma treated in a PelcoeasiGlow^TM^ discharge cleaning system at 0.39 mBar, 15 mA, for 30 s, and used blot times of 3 s.

### Cryo-EM data collection

For the datasets of pritelivir-bound HSV H/P complex and HSV replication fork complex, we collected datasets on a Cs-corrected Titan Krios (Thermo Fisher Scientific) operating at 300 kV, with post-GIF energy filter (20 eV slit) and a Gatan K3 camera in counting mode at 105,000x magnification, corresponding to a calibrated pixel size of 0.83 Å/pixel.

For the dataset of DNA-bound HSV H/P complex with pritelivir, 14,310 micrographs were collected at a dose rate of 16.65 e^−^ pixel^−1^ s^−1^. The total exposure time of 2.07 s was divided into 46 frames (total dose of approximately 50 e^-^/Å^2^).

For the dataset of DNA bound H/P complex with IM-250 or with amenamevir, we collected datasets on a Cs-corrected Titan Krios (Thermo Fisher Scientific) operating at 300 kV, with post-GIF energy filter (20 eV slit) and a Falcon 4 camera in counting mode at 165,000x magnification, corresponding to a calibrated pixel size of 0.73 Å/pixel. For the dataset of DNA-bound H/P complex with IM-250, 16,869 micrographs were collected at a dose rate of 8.9 e^−^ pixel^−1^ s^−1^. The total exposure time of 3 s was divided into 56 frames (total dose of approximately 52 e^-^/Å^2^). For the dataset of DNA bound H/P complex with amenamevir, 37,096 micrographs were collected at a dose rate of 11 e^−^ pixel^−1^ s^−1^. The total exposure time of 2.5 s was divided into 56 frames (total dose of approximately 51 e^-^/Å^2^). For the dataset of HSV replication fork complex, 25,091 micrographs were collected at a dose rate of 17.1 e^−^ pixel^−1^ s^−1^. The total exposure time of 2.07 s was divided into 46 frames (total dose of approximately 51 e^-^/Å^2^). A defocus range of -1.0 µm to -2.5 µm was used in all cases. The objective and C2 apertures were set to 100 µm and 50 µm, respectively.

### Cryo-EM image processing

We performed all image processing using RELION 3.0^63^ and cryoSPARC^64^. Movie frames were gain-normalized and motion-corrected using MotionCor2 1.6.4^65^. Contrast transfer function (CTF) correction was performed using CTFFind4.1^66^, as implemented in RELION 3.0.

For the pritelivir-bound H/P complex, we performed automated particle-picking with 9,118,480 particles (Figure S1b). After several rounds of 2D classifications, a subset of particles (128,342 in total) generated from the first round of heterogeneous refinement of 709,827 particles was subjected to further heterogeneous refinement, and a subset of particles (64,227 in total) generated from the second round of heterogeneous refinement was subjected to homogeneous refinement, yielding a final map of 3.7 Å resolution (Figure S1c). To obtain higher resolution maps for pritelivir-bound H/P complex, we masked the helicase–primase module and UL8 module for further local refinement, yielding final maps of 3.4 Å and 3.3 Å, respectively, with improved density (Figures S1d and S1e).

For the IM-250-bound H/P complex, we performed automated particle-picking with 5,535,965 particles (Figure S4b). After several 2D classifications, a subset of particles (177,199 in total) generated from the first round of heterogeneous refinement of 787,483 particles was subjected to the homogeneous refinement, yielding a final map of 2.9 Å resolution (Figure S4c). To obtain higher resolution maps for IM-250-bound H/P complex, we masked the helicase-primase module for further local refinement, yielding the final maps of 3.2 Å with improved density (Figure S4d).

For the amenamevir-bound H/P complex, automated particle-picking yielded 14,010,510 particles (Figure S5a). After several rounds of 2D classifications, a subset of 509,421 particles was selected for heterogeneous refinement, resulting in 340,875 particles used for homogeneous refinement, yielding a final map of 2.8 Å resolution (Figure S5b). To achieve higher-resolution maps for the amenamevir-bound H/P complex, we applied masks to the helicase-primase module and the UL8 module for further local refinement, yielding final maps at 2.6 Å resolution for both modules, each with improved density (Figures S5c and S5d).

For the DNA- and pritelivir-bound HSV replication fork complex, automated particle picking yielded 7,838,417 particles (Figure S11e). After several rounds of 2D classification, a subset of 1,652,862 particles was selected for a first round of heterogeneous refinement, resulting in 155,238 particles. These were subjected to a second round of heterogeneous refinement, yielding 60,225 particles for homogeneous refinement. This process produced a final map at 3.8 Å resolution (Figure S11f). To obtain higher-resolution maps for the HSV replication fork complex, focused masks were applied to the helicase-primase module and the polymerase module for local refinement, yielding final maps at 3.5 Å and 3.6 Å resolution, respectively, both with improved density (Figures S11g and S11h). Additionally, to improve resolution at the HSV replication fork complex interface, we applied a mask to the helicase-polymerase module and performed local refinement, yielding final maps at 3.7 Å and 4.1 Å with improved density, respectively (Figures S11i and S11j).

### Model building, refinement, and figure generation

For high-resolution cryo-EM structures of HPI-bound HSV H/P complex, we used an AF3 prediction^40^ of the HSV H/P complex as an initial model. To build the atomic models of the HPIs (pritelivir, IM-250, and amenamevir), the structures of HPIs were generated based on their SMILES formulas and fitted into the cryo-EM density in *Coot*^67^. The density corresponding to the HPI was clear (Figures S1f, S4e and S5e). We could only resolve the cryo-EM density of a 6 nt oligo in the helicase UL5. The rest of DNA segments were not visualized, likely because of flexibility.

The crystal and cryo-EM structures of HSV pol (PDB ID: 2GV9)^17^, UL42 (PDB ID: 1DML)^24^, and HSV polymerase holoenzyme (PDB ID: 8V1Q)^26^ were fitted as rigid bodies into the cryo-EM map using Chimera 1.15^68^. In the HSV replication fork complex, the density of UL42 and UL8 was weak, so we used the crystal and cryo-EM structures of UL42^24,26^ and the high-resolution model of UL8 from the HPI-bound H/P complex for docking into the density. We performed manual adjustments and iterative model building and real space refinement using *Coot* 0.9^67^ and PHENIX 1.20^69^. Figures were generated using PyMol v2.5.5 and UCSF ChimeraX v1.9^68^. For the HSV replication fork complex, we only saw poor density of the parental fork dsDNA at a low map contour level (see Figure S12a). We placed the dsDNA model into the density of parental DNA for figure generation but did not deposit it into final coordinates. MolProbity 4.5.2^70^ was used to validate the model. Statistics are provided in Table S3.

### Sequence alignments

The following sequences were used to generate alignments: HSV-1, herpes simplex virus 1 (GenBank: NC_001806.2); HSV-2, herpes simplex virus 2 (GenBank: NC_001798.1); VZV, varicella-zoster virus (GenBank: NC_001348.1); HCMV, human cytomegalovirus (GenBank: NC_006273.2); EBV, Epstein–Barr virus (GenBank: NC_007605.1); HHV-6A, human herpesvirus 6A (GenBank: NC_001664.2); KSHV, Kaposi’s sarcoma-associated herpesvirus (GenBank: NC_009333.1). Alignments were generated using Clustal Omega^71^ and plotted using ESPript 3.0.^72^

### AlphaFold 3 modeling

The predicted structure for the ATP-bound HSV H/P complex (UL5/UL52/UL8 from GenBank:NC_001806.2) was generated using AF3^40^. Modeling was performed using the 6-nt ssDNA (5′-TTTTTT-3′) along with one ATP molecule, one magnesium ion, and one zinc ion. The predicted FYNPYL motif interfaces were generated by providing residues comprising the motif for each virus and the corresponding H/P complex sequences. The following sequences were used for AF3^40^ modeling: HSV-1, GenBank: NC_001806.2; VZV, GenBank:NC_001348.1; HCMV, GenBank: NC_006273.2; HHV-6A, GenBank: NC_001664.2; EBV, GenBank: NC_007605.1; KSHV, GenBank: NC_009333.1.

### Mass photometry analysis of HSV replication fork complex

Mass photometry analyses were carried out with a Refeyn Two^MP^ mass photometer (Refeyn Ltd, Oxford, UK) at room temperature. Glass coverslips and gaskets were cleaned with HPLC-grade water and isopropanol and dried under filtered gas before use. HSV H/P complex alone (UL5/UL52/UL8), polymerase holoenzyme alone (UL30/UL42), and replication fork complex components (UL5/UL52/UL8+UL30/UL42) with or without DNA substrate were diluted to 200 nM in buffer containing 25 mM HEPES-NaOH, pH 7.5, 150 mM NaCl, 2 mM MgCl_2_ and 1 mM DTT. Eighteen microliters of buffer were used to find the camera focus prior to loading 2 μl of the sample onto the gasket. Acquisition camera image size was set to medium. Data were collected as a 1 min movie and then processed using ratiometric imaging. To correlate ratiometric contrast to mass, the Refeyn Two^MP^ instrument was calibrated using molecular standards of monomeric bovine serum albumin (BSA) (66 kDa), dimeric BSA (132 kDa), and thyroglobulin (MW 660 kDa), with a molecular weight error of less than 5%. The experiment was performed twice. Data were analyzed using Discover^MP^ version 2.3 software (Refeyn Ltd, Oxford, UK).

### MD simulations

The cryo-EM structures of the HSV-DNA complex with pritelivir or amenamevir were prepared prior to modelling and simulations. The Protein Preparation module in Schrödinger Maestro 2025-1^73^ was applied to cap termini, repair residues, optimize H-bond assignments, and run restrained minimizations using default settings. Protein interactions with ligand, categorized into five groups: hydrogen bonds, hydrophobic, ionic, water bridges, and halogen bonds, were monitored throughout the simulation using Schrödinger’s interaction diagram. The interaction fraction of each interaction type, normalized against the total frames of the simulation, was plotted.

### Rolling-circle assay

To determine the helicase activity of HSV UL5 and the polymerase activity of HSV UL30, we performed a helicase assay. The 90 nt oligonucleotides 5′ 6-FAM-labeled 5′-TTTTTTTTTTTTTTTTTTTTTTTTTTTTTTTTTTTTTTTTTTTTTTTTTTTTTTTTTTTTTTTTCCCAGTCACGACGTTGTAAAACGACGGCCAGT-3′ with 60 nt mismatch poly T, and M13mp18 ssDNA (NEB Cat# N4040S) were annealed to generate the ssDNA vector with a fork (Table S2). Rolling-circle assays with 6-FAM-labeled substrates (50 nM), HSV polymerase holoenzyme (0.4 μM), and H/P complex (0.4, 0.8, 1.6, and 2.4 μM), and 0.8 μM ICP8 were performed in 25 mM HEPES pH 7.5, 150 mM NaCl, 10 mM MgCl_2_, 10 mM ATP,1 mM dNTP and 0.1 mg ml^-1^ BSA (Sigma-Aldrich Cat# A7906). The reaction mixture was incubated at 4°C for 10 min and then incubated at 37 °C for 2 h. To terminate all reactions, we added 20 mM EDTA, 0.5% (v/v) SDS (sodium dodecyl-sulfate) and 6X agarose gel loading dye (Thermo Fisher Scientific Cat# J62157). Products were separated on a 0.8% alkaline agarose gel, and gels were imaged with a Typhoon FLA 9500 (GE Healthcare), with signals quantified using Image Studio (LI-COR) v5.2. To quantify the DNA product levels, a box of equivalent size was used for other bands of the same M13 ssDNA samples was drawn at the corresponding position on the left negative control (NC, -/-) lane and used to set the background to 0. The resulting values were then normalized to the corresponding value from the reaction with 0.4 μM polymerase holoenzyme and 2.4 μM H/P complex, which was set to 1.

For testing the inhibition effects of HPIs, rolling-circle assays with 6-FAM-labeled substrates (50 nM), HSV polymerase holoenzyme (0.4 μM), H/P complex (0.4, 0.8, 1.6, and 2.4 μM), ICP8 (0.8 μM), and HPIs (pritelivir, amenamevir) (0.1, 1, and 10 μM) were performed in 25 mM HEPES pH 7.5, 150 mM NaCl, 10 mM MgCl_2_, 10 mM ATP,1 mM dNTP and 0.1 mg ml^-1^ BSA.

Reactions were incubated at 4 °C for 10 min and then incubated at 37 °C for 2 h. We added 20 mM EDTA, 0.5% (v/v) SDS and 6X agarose gel loading dye (Thermo Cat#J62157) to terminate the reactions. Products were separated on a 0.8% alkaline agarose gel, and gels were imaged with Typhoon FLA 9500 (GE Healthcare), with signals were quantified using Image Studio (LI-COR) v5.2. To quantify the DNA product levels, a box of equivalent size as used for other bands of the same M13 ssDNA samples was drawn at the corresponding position on the NC lane and used to set the background to 0. The resulting values were then normalized to the corresponding value from the positive control (PC) reaction, which was set to 1.

### Optical tweezer experiments

Single-molecule experiments were performed on a C-Trap (LUMICKS) integrating optical tweezers and microfluidics. The five-channel laminar flow cell was used for experiments after passivation using 0.5% (w/v) Pluronic F128 in phosphate buffered saline (PBS), and subsequently with BSA (1 mg ml^−1^).

We performed the experiment with WT H/P complex or H/P complex with UL5 K356N or UL52 F360R mutant subunits. Streptavidin-coated polystyrene beads (4.35 µm, Spherotech Cat# SVP-40-5) at a concentration of 0.005% (w/v) were injected into channel 1. A biotin-labeled 17 kb DNA molecule, which contained two nicks (5 kb apart) on one of the strands (about 2 pM) (LUMICKS Cat# 00027), was introduced into channel 2. Buffer A, containing 25 mM HEPES (pH 7.5), 150 mM NaCl, 0.1 mg ml^−1^ BSA, and 1 mM DTT, was injected into channel 3. The HSV H/P complex was diluted to 100 nM in Buffer A and introduced into channel 4. In parallel, a mixture of the H/P complex and pritelivir in Buffer A (the final concentration of H/P complex was 100 nM, and the final concentrations of pritelivir were 100, 200, and 500 nM) was introduced into channel 5. After completion of measurements, both channels were flushed with Buffer A for 10 min to remove residual protein or inhibitor. Subsequently, the next set of experimental conditions was applied to channels 4 and 5. This cycle was repeated until all experimental conditions had been tested.

The optical traps were calibrated using the power spectrum of Brownian motion of the trapped beads to achieve a trap stiffness of 0.16–0.18 pN nm^−1^. After optically trapping two beads, the DNA molecule was tethered between the beads under flow in channel 2. The presence of a single DNA tether was verified by measuring a force–extension curve at a constant pulling rate of 0.2 µm s^−1^ and comparing it to the worm-like chain model of dsDNA. The tethered DNA molecule was extended beyond the contour length of the DNA (6.34 μm) to around 8.5 μm and held for 10 s in the presence of flow to melt away a piece of ssDNA leaving behind a gap of 5 kb on the tethered DNA molecule. Subsequently, the DNA tether was moved to the protein channel (channels 4 and 5) and incubated for 10–30 s. In most of the experiments, this loading step was performed using DNA held at a very low force (∼1 pN). After protein loading, the unwinding experiment was performed in the same channel at constant force (45 pN). The changes in distance between beads were recorded for further analysis. All experiments were performed at room temperature (28 °C).

### Optical tweezer data analysis

All data analysis was carried out in Python using custom-written scripts. Helicase activity, recorded as the change in distance between optically trapped beads, was converted to the number of nucleotides unwound^74^.

From each raw trace, multiple 15 s windows (approximately 250 data points per window) were extracted. These segments were smoothed using a Savitzky–Golay filter, after which a linear regression was performed on each window to estimate the unwinding rates.

To identify pause states, the raw data were first filtered using the Savitzky-Golay filter to reduce noise. The discrete derivative of the unwound nucleotides with respect to time was then computed. A threshold of 2 nucleotides per second (nt/s) was used; time points with instantaneous unwinding rates below this threshold were classified as ‘paused’. The fraction of paused states was quantified using 50 s sliding windows across the filtered trace (Figure S10c).

Statistical comparisons between experimental conditions were performed using Welch’s t-test. Pairwise comparisons were conducted as indicated in the text. All statistical analyses were performed in Python using standard scientific libraries.

---

**Figure S1.**
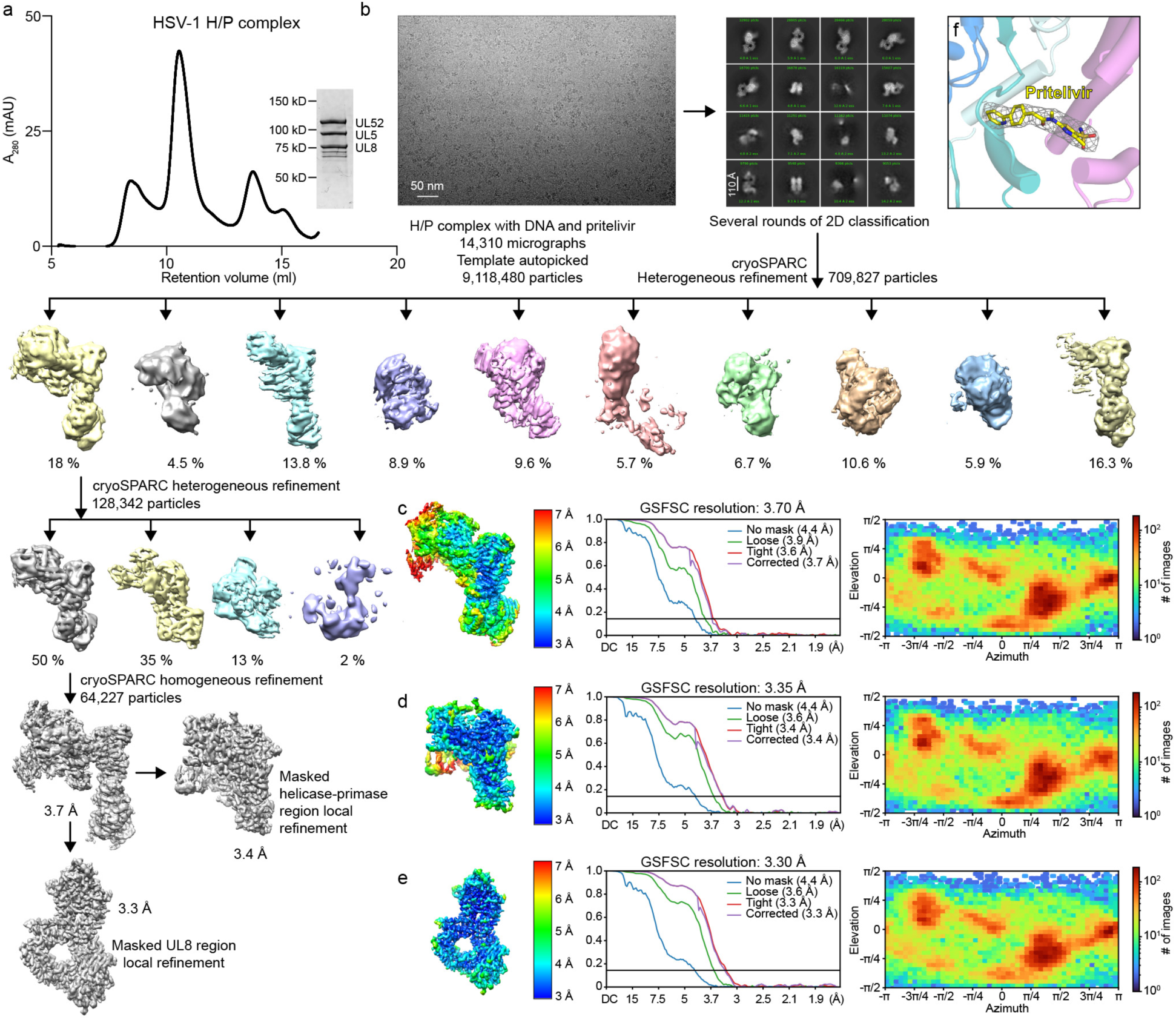
Structure determination of the pritelivir-bound HSV H/P complex. (a) Size-exclusion chromatography profile and SDS-PAGE analysis of the HSV H/P complex using a Superdex 200 Increase column. The retention volume of the main peak is indicated for each chromatogram. The bands that are smaller than UL8 likely represent degradation products or contaminants. (b) Workflow used for cryo-EM data processing of pritelivir-bound HSV H/P complex. (c–e) Local resolution estimation, Fourier shell correlation (FSC) curves, and particle angular distributions of the cryo-EM reconstructions of the overall pritelivir-bound HSV H/P complex at 3.7 Å resolution (c), masked helicase-primase region at 3.4 Å resolution (d) and masked UL8 region at 3.3 Å resolution (e). (f) Cryo-EM density of pritelivir in the drug binding site of the HSV H/P complex. Pritelivir is shown as sticks.

**Figure S2.**
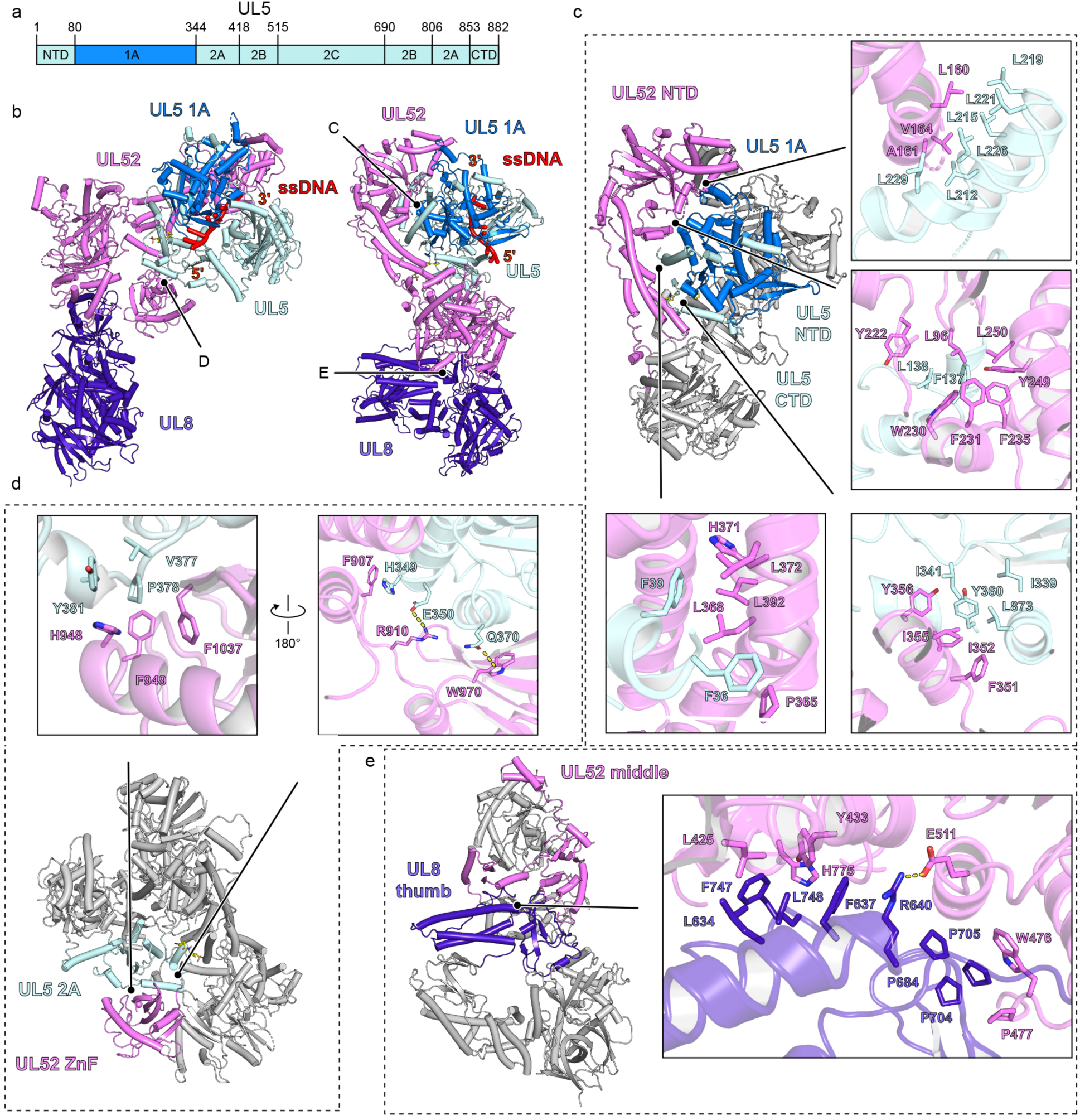
Domain architecture of the HSV H/P complex. (a) Schematic diagram of the domain architecture of the HSV helicase UL5 and primase UL52. (b) Ribbon diagram of the HSV H/P complex bound to ssDNA and pritelivir. (c) Interactions between the primase UL52 NTD and UL5 NTD, 1A domain, and CTD. (d) Interactions between the primase UL52 ZnF domain and UL5 2A domain. (e) Interactions between the primase UL52 middle domain and UL8 thumb domain.

**Figure S3.**
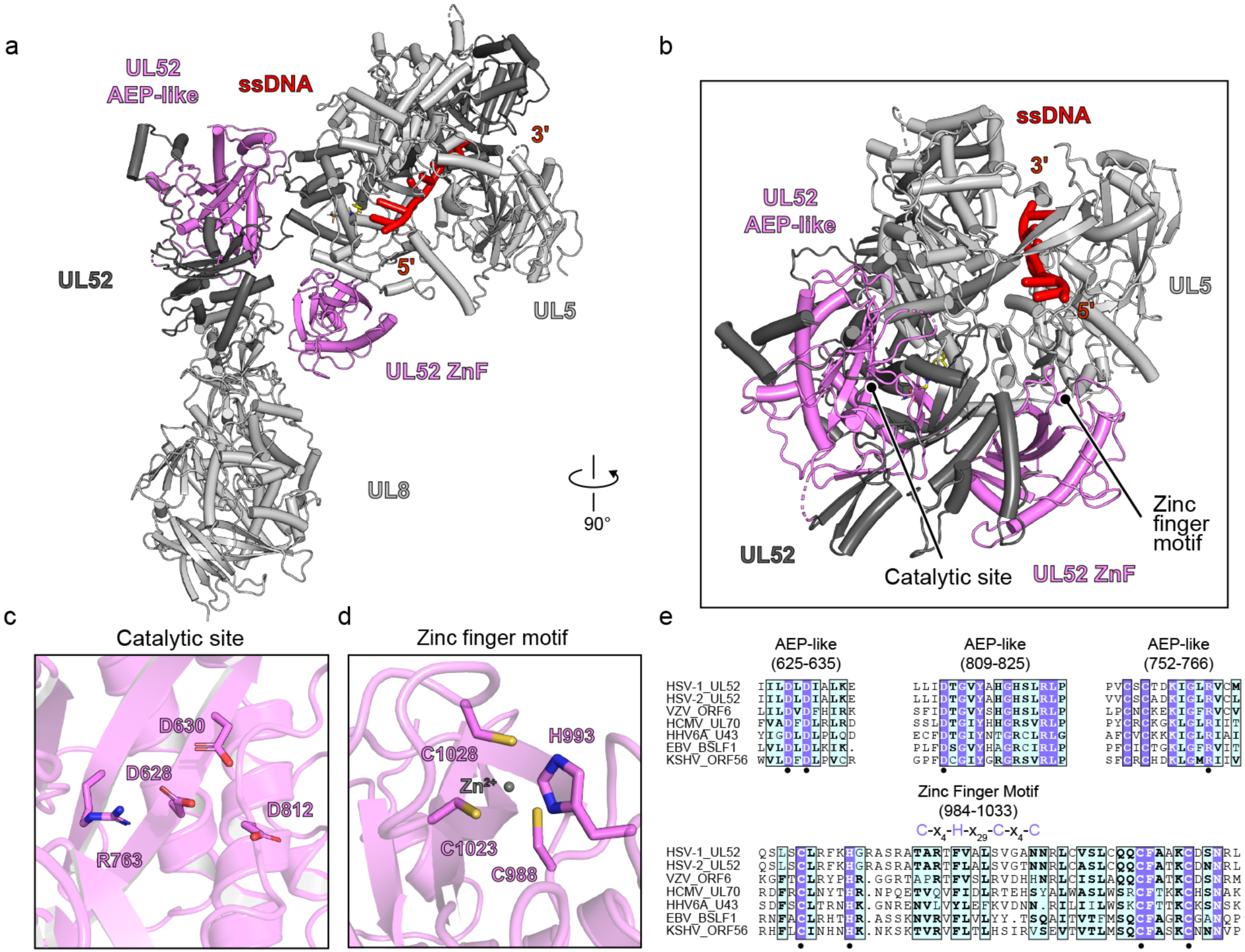
The HSV H/P complex primase active site. (a) Ribbon diagram of the HSV H/P complex bound to ssDNA and pritelivir. The AEP-like and ZnF domains in UL52 are labeled. (b) Close-up view of the UL52 AEP-like and ZnF domains. The locations of the AEP-like active site and zinc finger motifs are indicated. (c) HSV primase UL52 catalytic site in the AEP-like domain. (d) HSV primase UL52 zinc binding site in the ZnF domain. A zinc ion is shown as a sphere. (e) Sequence alignment of herpesvirus primase active sites and zinc finger motifs. Conserved active site residues are indicated. HSV-1, herpes simplex virus 1; HSV-2, herpes simplex virus 2; VZV, varicella-zoster virus; HCMV, human cytomegalovirus; EBV, Epstein–Barr virus; HHV-6A, human herpesvirus 6A. KSHV, Kaposi’s sarcoma-associated herpesvirus.

**Figure S4.**
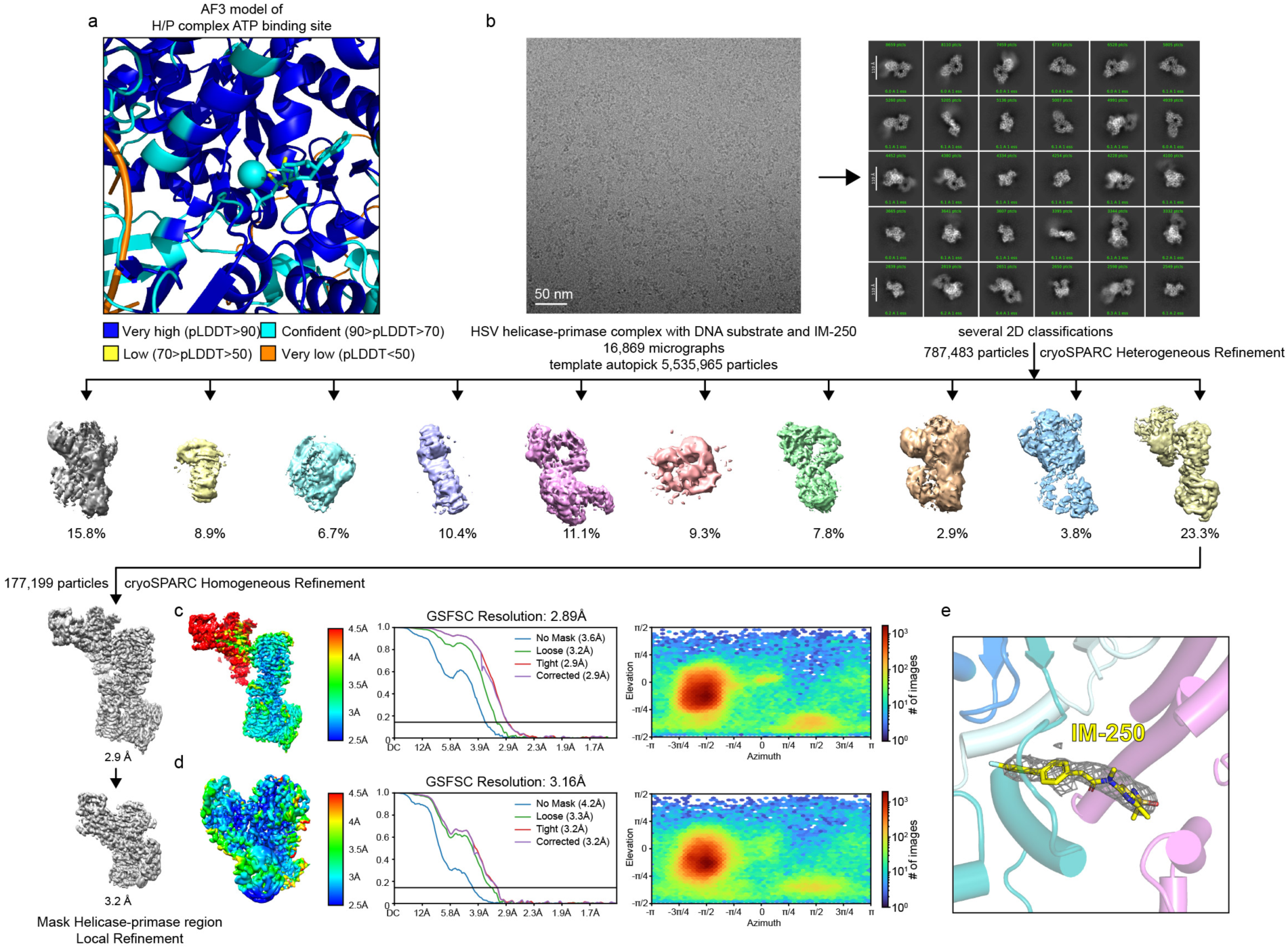
Structure determination of the IM-250 bound HSV H/P complex. (a) pLDDT of the ATP binding site of the AF3 model of ATP-bound HSV H/P complex. (b) Workflow used for cryo-EM data processing of IM-250-bound HSV H/P complex. (c and d) Local resolution estimation, Fourier shell correlation (FSC) curves, and particle angular distributions of the cryo-EM reconstructions of the IM-250-bound HSV H/P complex at 2.9 Å resolution (c) and masked helicase–-primase region at 3.2 Å resolution (d). (e) Cryo-EM density of IM-250 in the drug binding site of the HSV H/P complex. IM-250 is shown as sticks.

**Figure S5.**
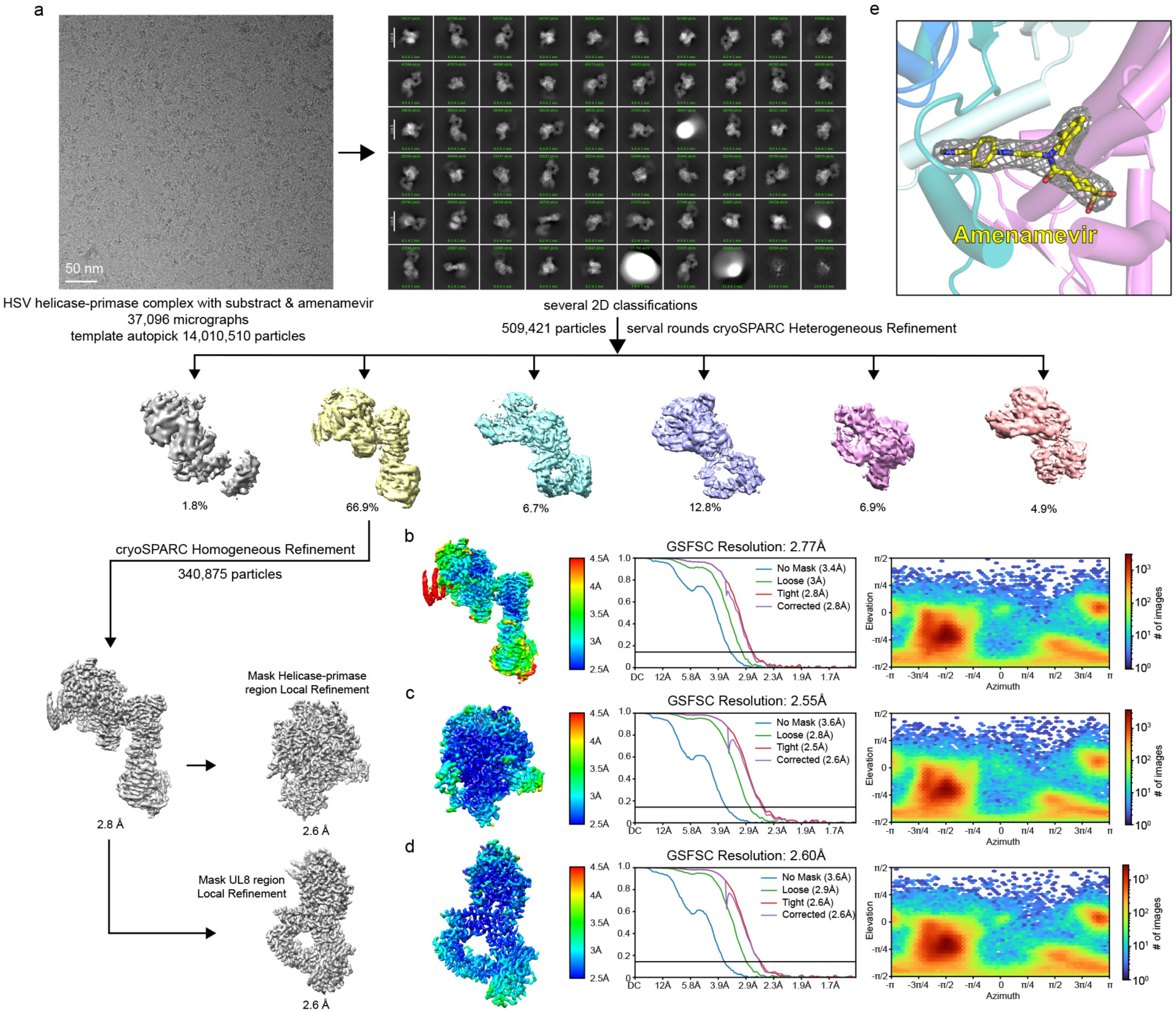
Structure determination of the amenamevir bound HSV H/P complex. (a) Workflow used for cryo-EM data processing of amenamevir-bound HSV H/P complex. (b–d) Local resolution estimation, Fourier shell correlation (FSC) curves, and particle angular distributions of the cryo-EM reconstructions of the amenamevir-bound HSV H/P complex at 2.8 Å resolution (b), the masked helicase–primase region at 2.6 Å resolution (c), and the masked UL8 region at 2.6 Å resolution (d). (e) Cryo-EM density of amenamevir in the drug binding site of the HSV H/P complex. Amenamevir is shown as sticks.

**Figure S6.**
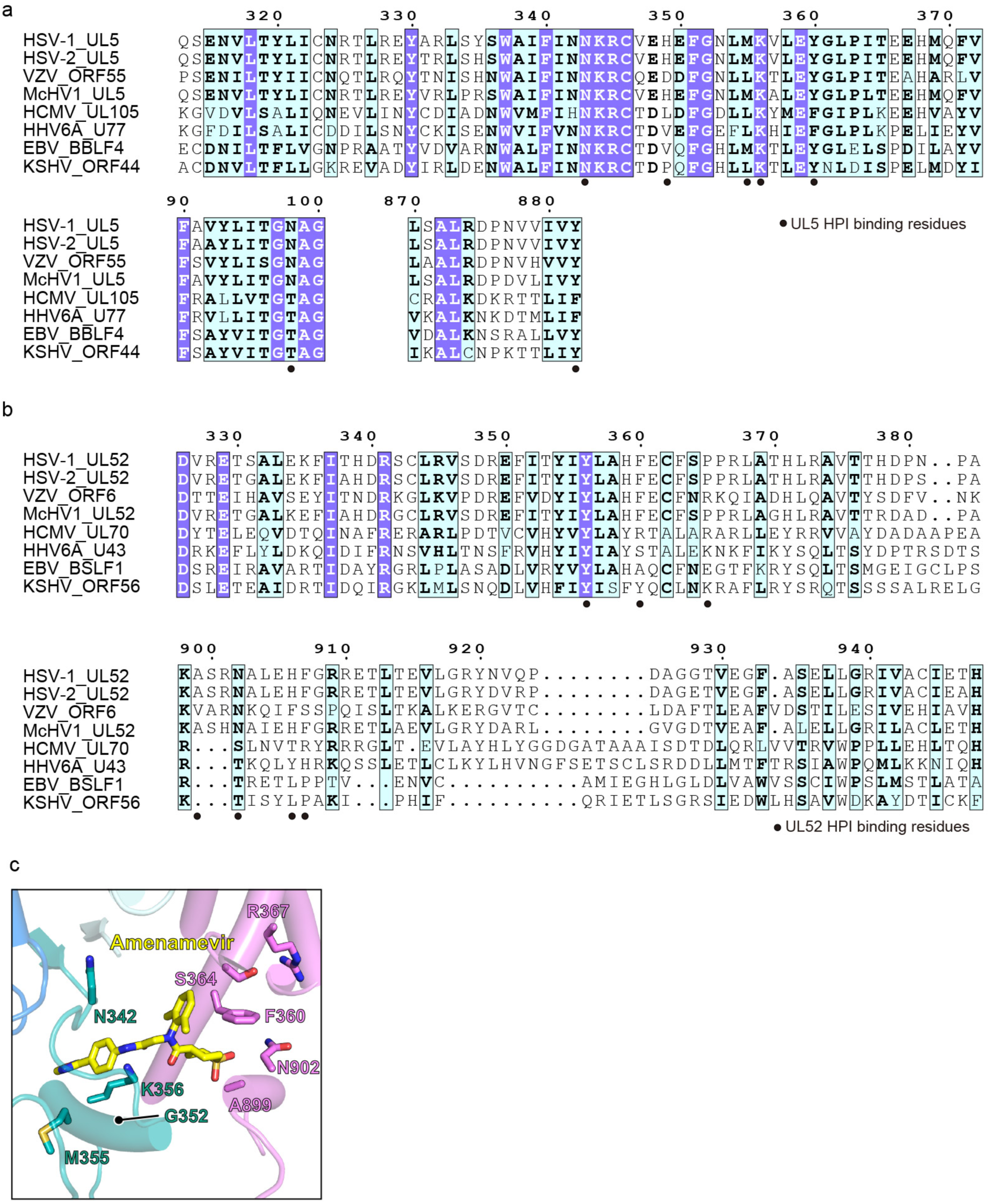
Sequence alignment showing UL5 and UL52 residues that interact with HPIs. (a and b) Sequence alignments of UL5 (a) or UL52 (b) regions that contain residues interacting with HPIs. These residues are indicated with circles. HSV-1, herpes simplex virus 1, herpes simplex virus 2; VZV, varicella-zoster virus; McHV1, macacine alphaherpesvirus 1; HCMV, human cytomegalovirus; EBV, Epstein–Barr virus; HHV-6A, human herpesvirus 6A; KSHV, Kaposi’s sarcoma-associated herpesvirus. (c) Close-up view of the amenamevir binding site. Residues that are sites of HPI resistance mutations are shown. See Table S5 for additional information.

**Figure S7.**
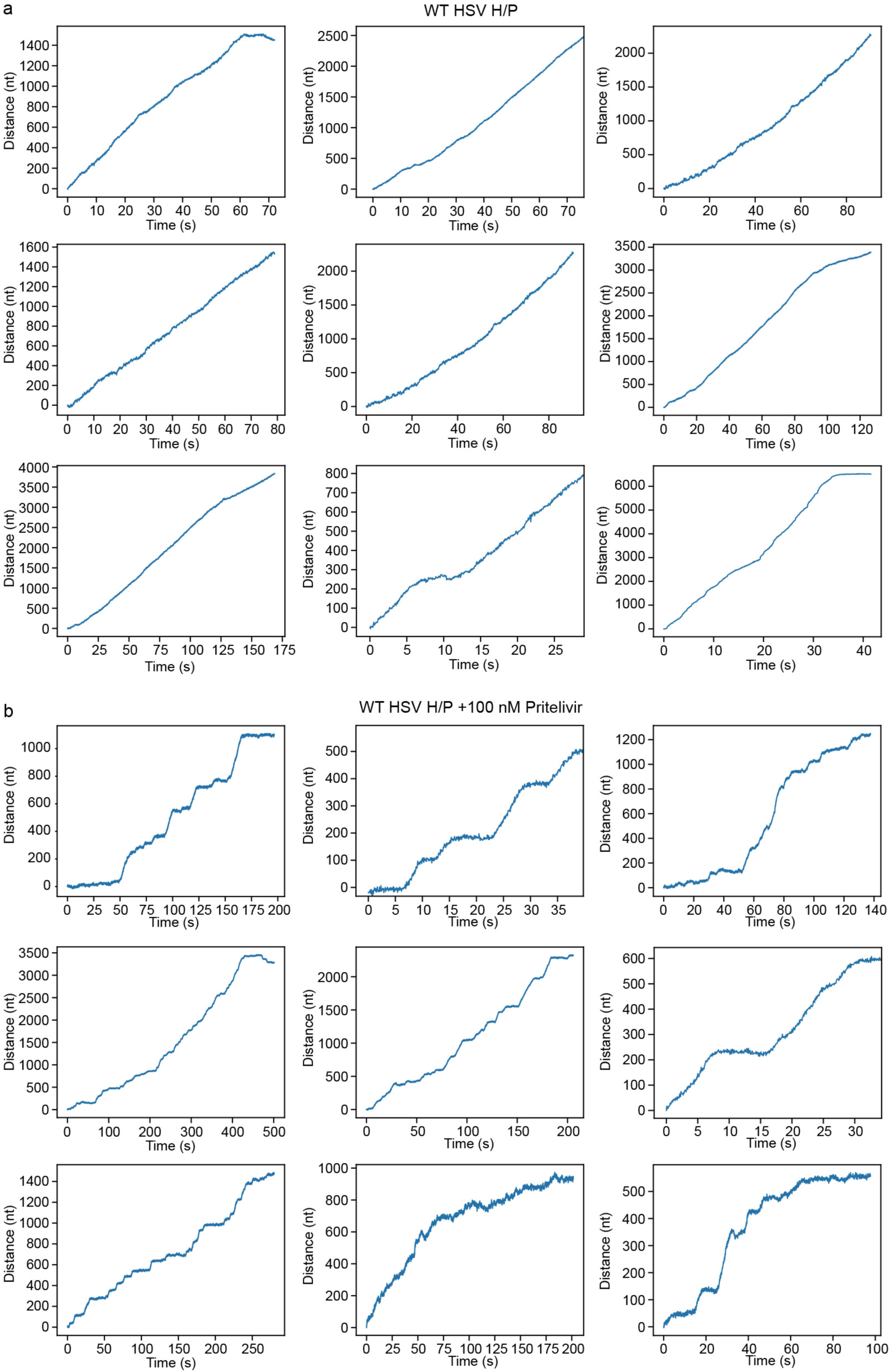
Representative optical tweezer traces of WT HSV H/P complex with and without pritelivir. (a) Traces of unwinding activity in the presence of 100 nM HSV H/P complex measured in optical tweezer experiments. The experiments with 100 nM H/P complex alone were repeated 16 times. (b) Traces of unwinding activity in the presence of 100 nM HSV H/P complex and 100 nM pritelivir measured in optical tweezer experiments. The experiments with H/P complex and 100 nM pritelivir were repeated 11 times.

**Figure S8.**
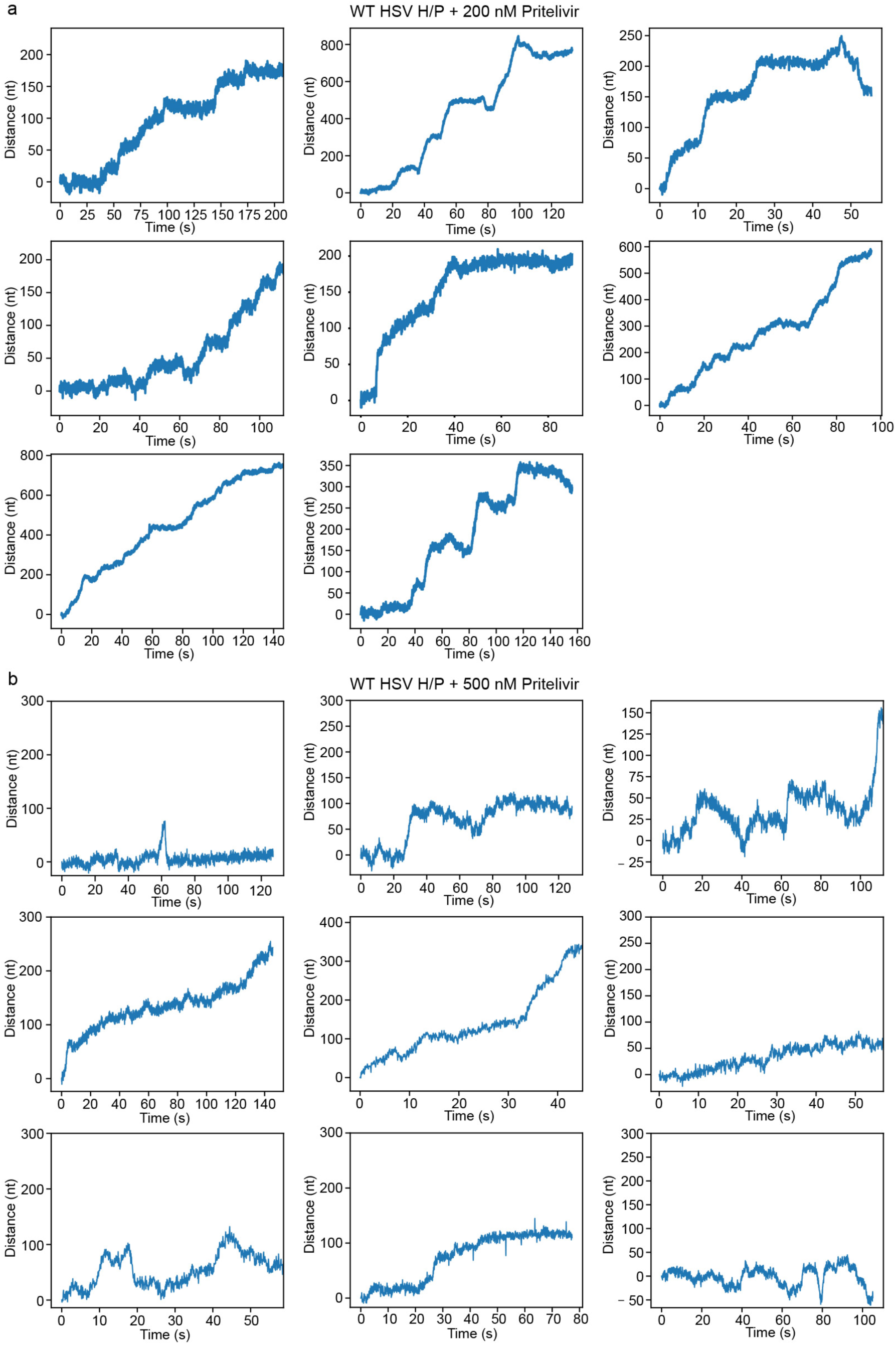
Representative optical tweezer traces of WT HSV H/P complex with pritelivir. (a) Traces of unwinding activity in the presence of 100 nM HSV H/P complex and 200 nM pritelivir measured in optical tweezer experiments. The experiments with H/P complex and 200 nM pritelivir were repeated 18 times. (b) Traces of unwinding activity in the presence of 100 nM HSV H/P complex and 500 nM pritelivir measured in optical tweezer experiments. The experiments of H/P complex with 500 nM pritelivir were repeated 12 times.

**Figure S9.**
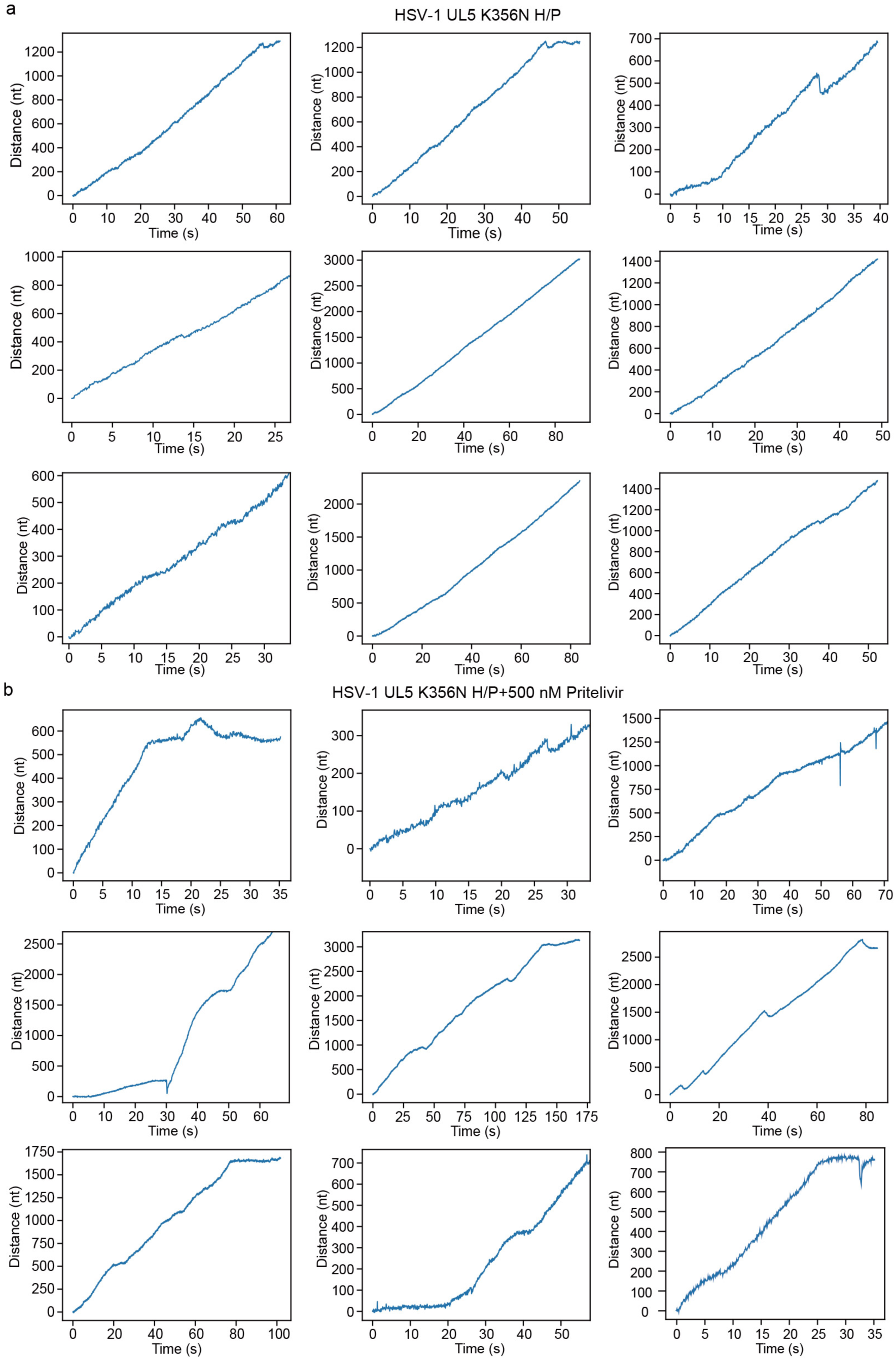
Representative optical tweezer traces of HSV UL5 K356N H/P complex with and without pritelivir. (a) Traces of unwinding activity in the presence of 100 nM HSV UL5 K356N H/P complex measured in optical tweezer experiments. (b) Traces of unwinding activity in the presence of 100 nM HSV UL5 K356N H/P complex and 500 nM pritelivir measured in optical tweezer experiments. The experiments of H/P complex with UL5 K356N with or without 500 nM pritelivir were repeated 15 and 17 times, respectively.

**Figure S10.**
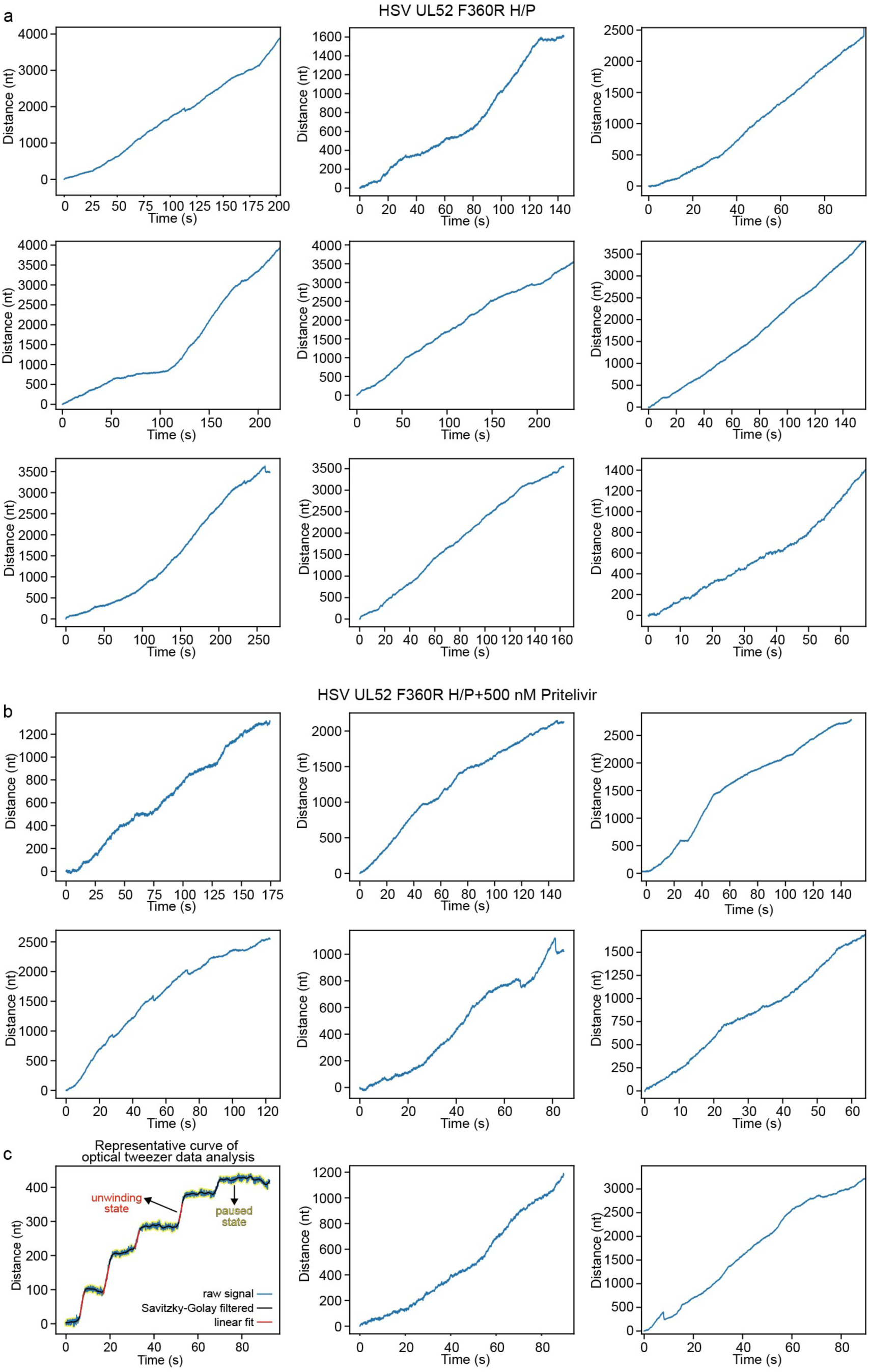
Representative optical tweezer traces of HSV UL52 F360R H/P complex with and without pritelivir. (a) Traces of unwinding activity in the presence of 100 nM HSV UL52 F360R H/P complex measured in optical tweezer experiments. (b) Traces of unwinding activity in the presence of 100 nM HSV UL52 F360R H/P complex and 500 nM pritelivir measured in optical tweezer experiments. The experiments of H/P complex with UL52 F360R with or without 500 nM pritelivir were repeated for 11 and 11 times, respectively. (c) Representative curve of optical tweezer data analysis. This curve is from WT HSV H/P complex with 200 nM pritelivir. The paused state and the unwinding state are labeled after classification based on instantaneous unwinding rate (see Methods).

**Figure S11.**
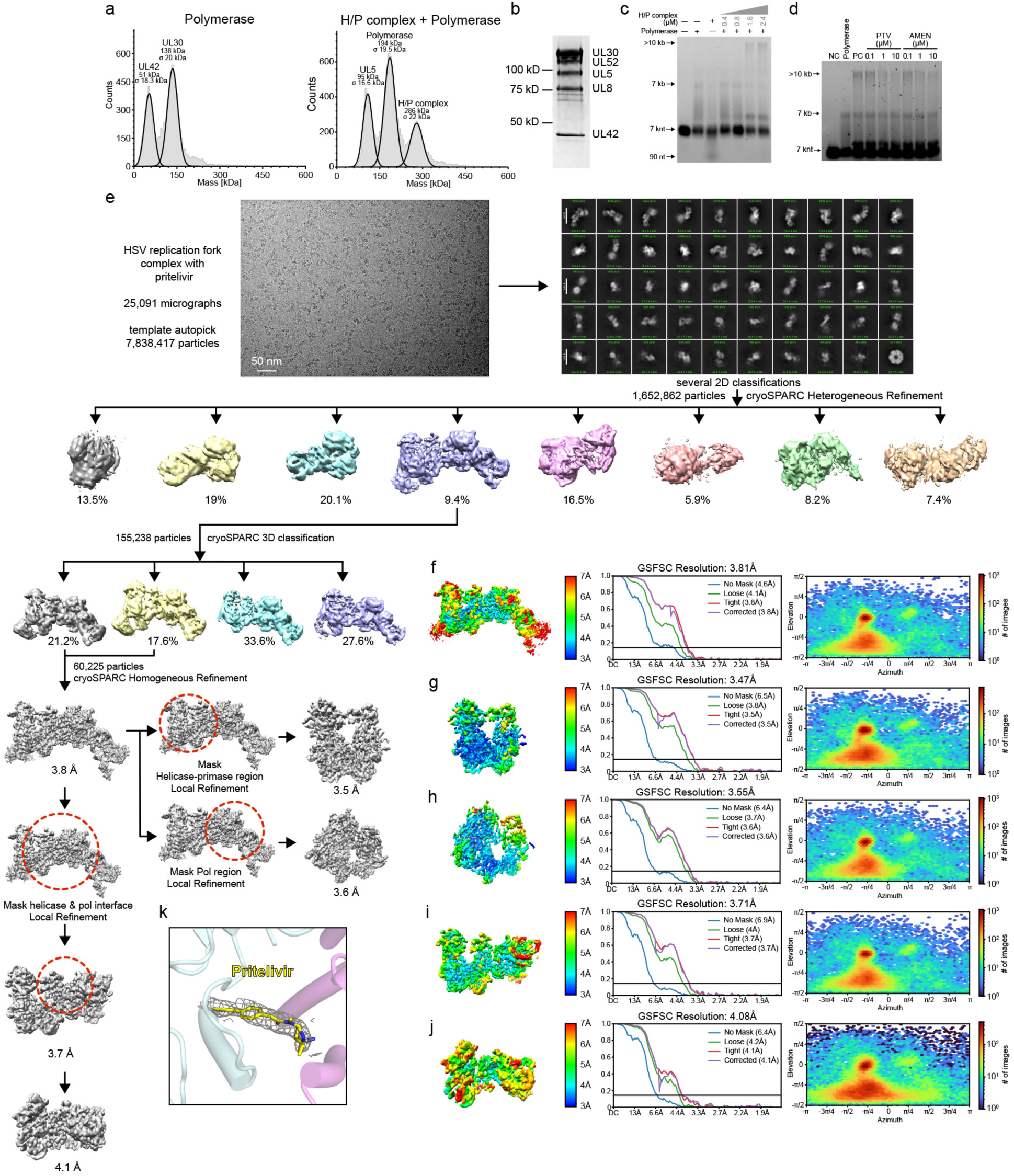
Mass photometry results, rolling-circle assay gels, and structure determination of the pritelivir-bound HSV replication fork complex. (a) Mass photometry analysis of the HSV polymerase holoenzyme (UL30/UL42) alone or the HSV replication fork complex components (UL5/UL52/UL8+ UL30/UL42) without DNA. The experiment was performed twice, and representative data are shown. (b) SDS–PAGE analysis of the HSV replication fork complex after size-exclusion chromatography, visualized using a stain-free gel system. (c) Products of the rolling circle assay performed with HSV polymerase with or without co-incubation with the H/P complex visualized using alkaline agarose gel electrophoresis. This experiment was performed three times, and a representative gel is shown. (d) Rolling-circle assay performed with the HSV replication fork complex co-incubated with different concentrations of pritelivir (PTV) or amenamevir (AMEN), visualized using alkaline agarose gel electrophoresis. This experiment was performed three times, and a representative gel is shown. (e) Workflow used for cryo-EM data processing of the HSV replication fork complex. (f–j) Local resolution estimation, Fourier shell correlation (FSC) curves, and particle angular distributions of the cryo-EM reconstructions of the overall HSV replication fork complex at 3.8 Å resolution (f), masked HSV helicase at 3.5 Å resolution (g), masked HSV polymerase at 3.6 Å resolution (h), masked interface between polymerase and helicase–primase complex at 3.7 Å resolution (i) and masked FYNPYL motif binding site at 4.1 Å resolution (j). (k) Cryo-EM density of pritelivir in the drug binding site in the HSV replication fork complex structure. Pritelivir is shown as sticks.

**Figure S12.**
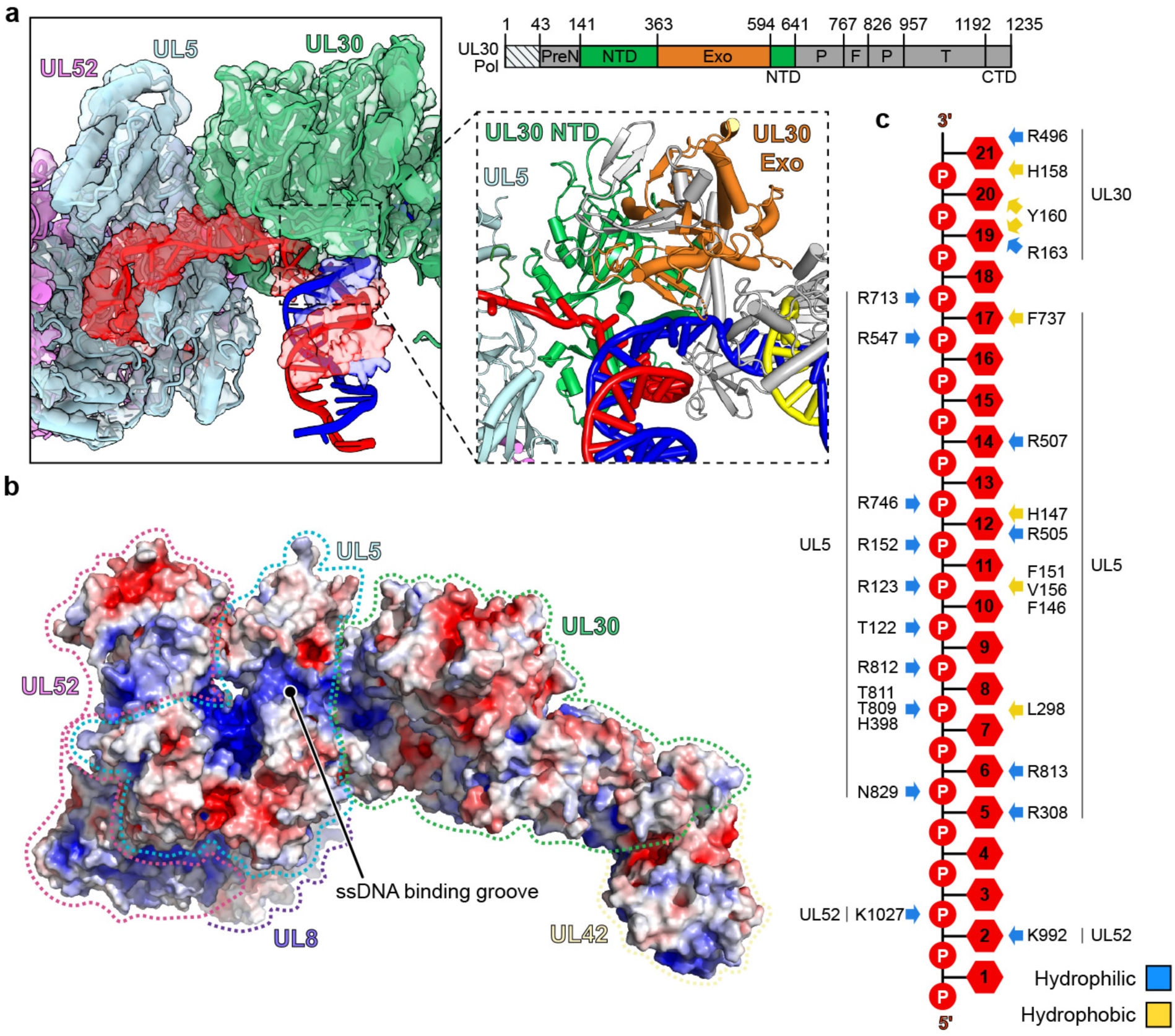
HSV replication fork complex ssDNA and parental dsDNA interactions. (a) Cryo-EM density (left) of parental DNA in the HSV replication fork complex shown as a surface. The presumed lagging strand is colored red. The presumed leading strand is colored blue. Close-up view (right) of the location of parental DNA. The UL30 NTD shown in green. The UL30 Exo domain is shown in orange. The remaining domains in UL30 are shown in light gray. (b) Electrostatic surface potential representation of the HSV replication fork complex ssDNA binding groove. Each subunit is labeled. (c) Schematic representation of the HSV replication fork complex interactions with the presumed lagging strand DNA. Hydrophilic interactions are shown in blue. Hydrophobic interactions are shown in yellow.

**Figure S13.**
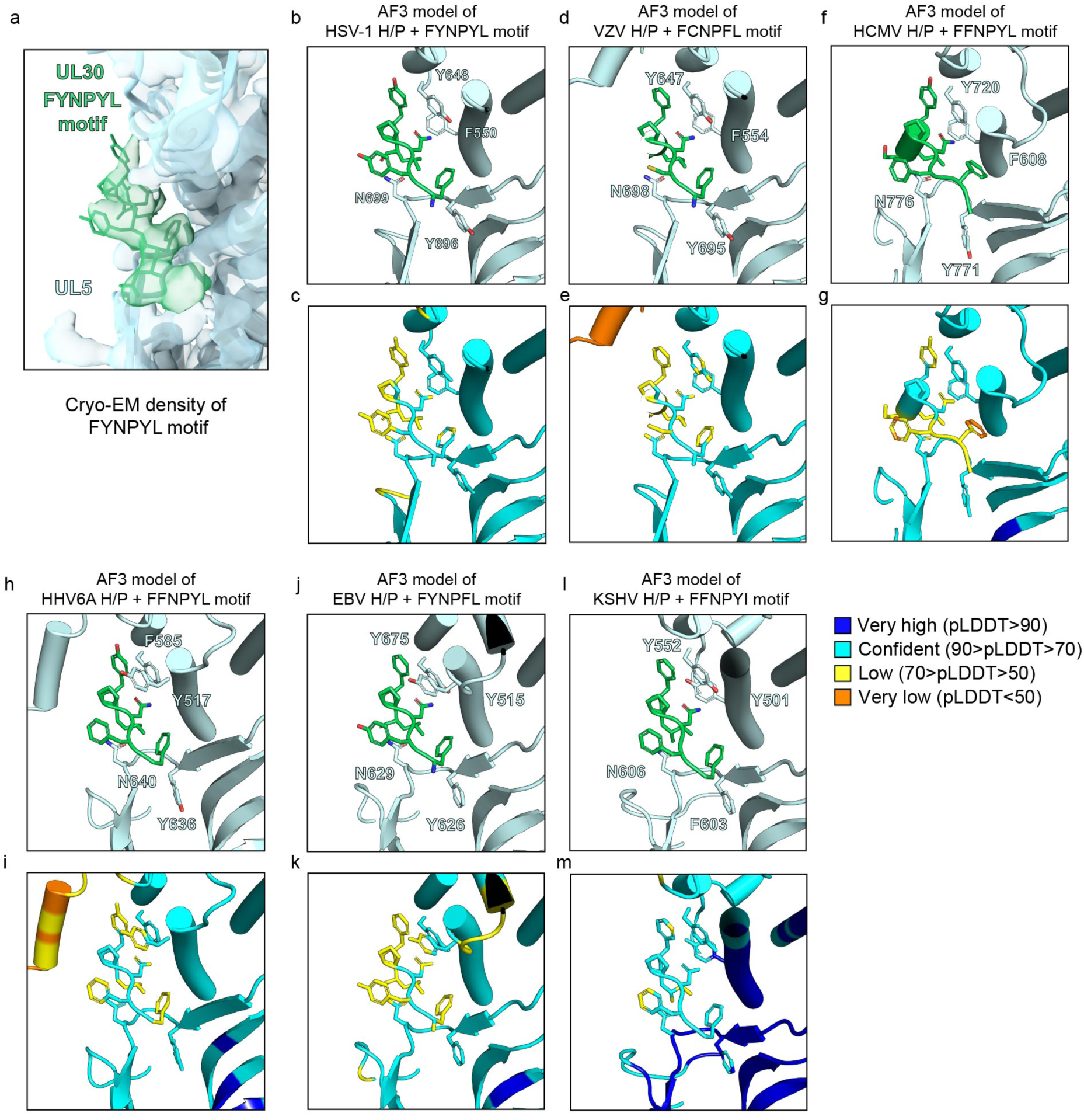
AF3 models of herpesvirus H/P complex bound to polymerase FYNPYL motif. (a) Cryo-EM density of the UL30 FYNPYL motif in the HSV replication fork complex shown as surface. The UL30 FYNPYL motif is shown as sticks (b–m) Structures and pLDDT scores of AF3 models of herpesvirus H/P complexes bound to the polymerase preN domain containing the indicated motifs: FYNPYL (HSV, b and c); FCNPFL (VZV, d and e); FFNPYL (HCMV, f and g); FFNPYL (HHV-6A, h and i); FYNPFL (EBV, j and k); and FFNPYI (KSHV, l and m).

**Figure S14.**
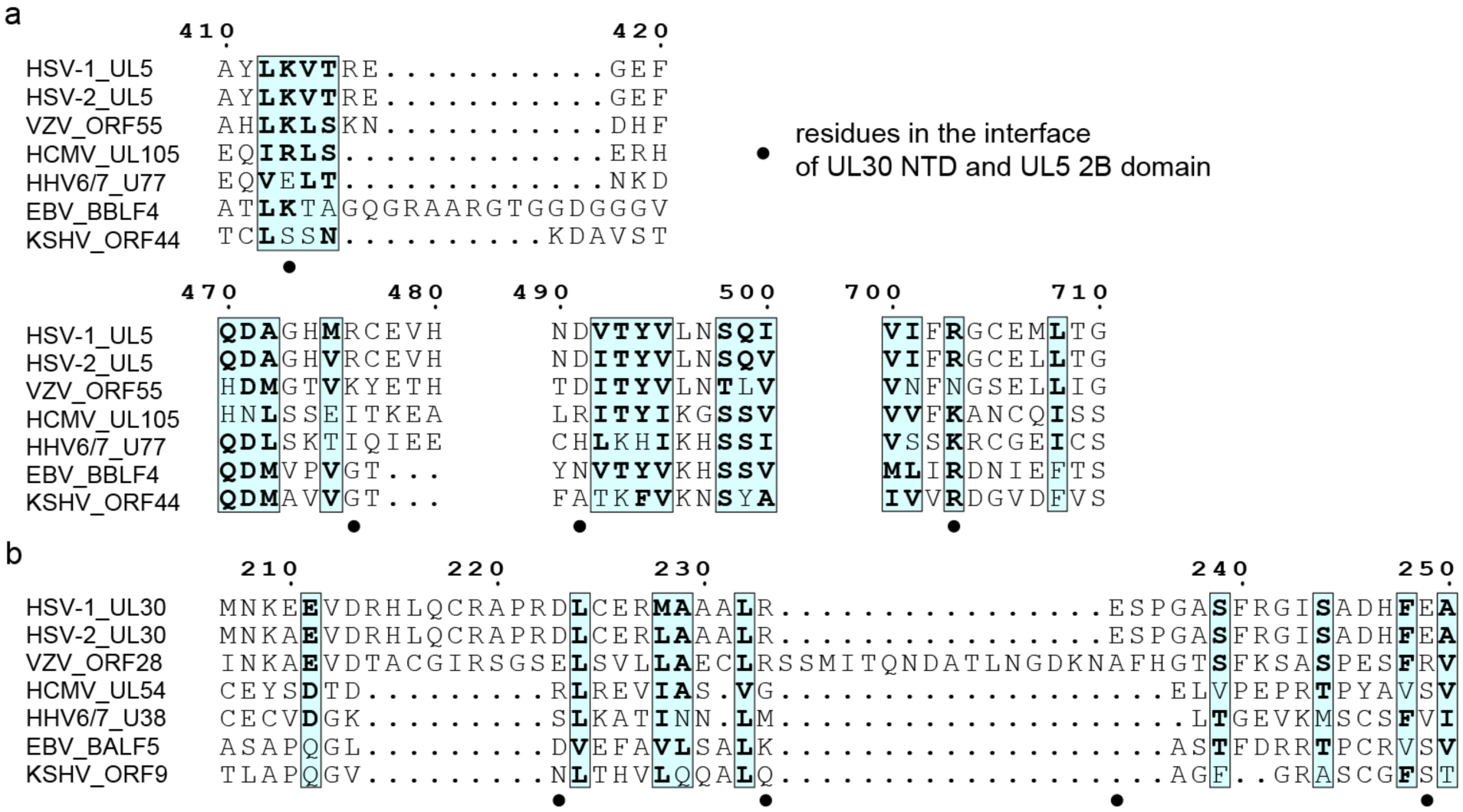
Sequence alignment of interacting residues at the UL5 2B–UL30 NTD interface. (a and b) Sequence alignment of the HSV UL5 2B domain (a) or the UL30 NTD (b) with the analogous domains of other herpesviruses. Residues involved in interactions at the interface of these domains are indicated. HSV-1, herpes simplex virus 1; HSV-2, herpes simplex virus 2; VZV, varicella-zoster virus; HCMV, human cytomegalovirus; EBV, Epstein–Barr virus; HHV-6A, human herpesvirus 6A; KSHV: Kaposi’s sarcoma-associated herpesvirus.

**Table S1.** Expected masses of subunits and complexes and mass photometry results.

| Complex | Subunit mass | Complex mass | Measured mass |
| --- | --- | --- | --- |
| Helicase–primase (UL5/UL52/UL8) | 99 kDa (UL5)<br>114 kDa (UL52)<br>80 kDa (UL8) | 293 kDa | 292 s.d. of ± 20 kDa |
| Polymerase holoenzyme (UL30/UL42) | 131 kDa (UL30)<br>35 kDa (UL42) | 166 kDa | 196 s.d. of ± 33 kDa |
| Replication fork complex (RFC) with DNA | 460 kDa (RFC)<br>30 kDa (DNA) | 490 kDa | 494 s.d. of ± 52 kDa |

**Table S2.**
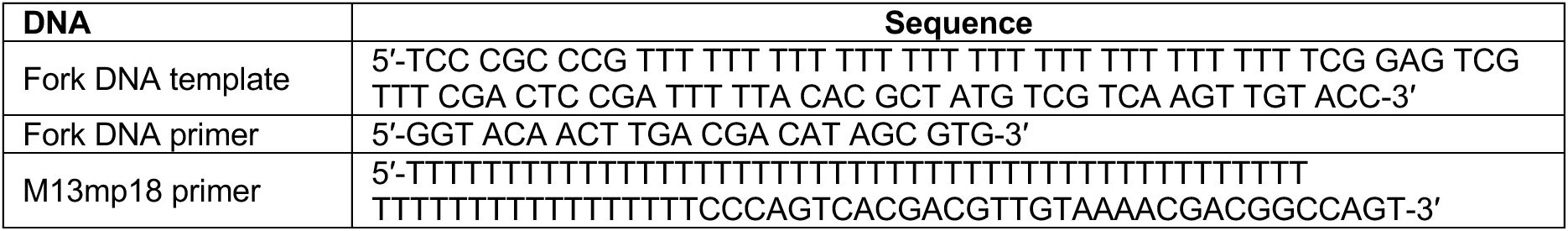
DNA substrates for cryo-EM and rolling-circle assays.

**Table S3.**
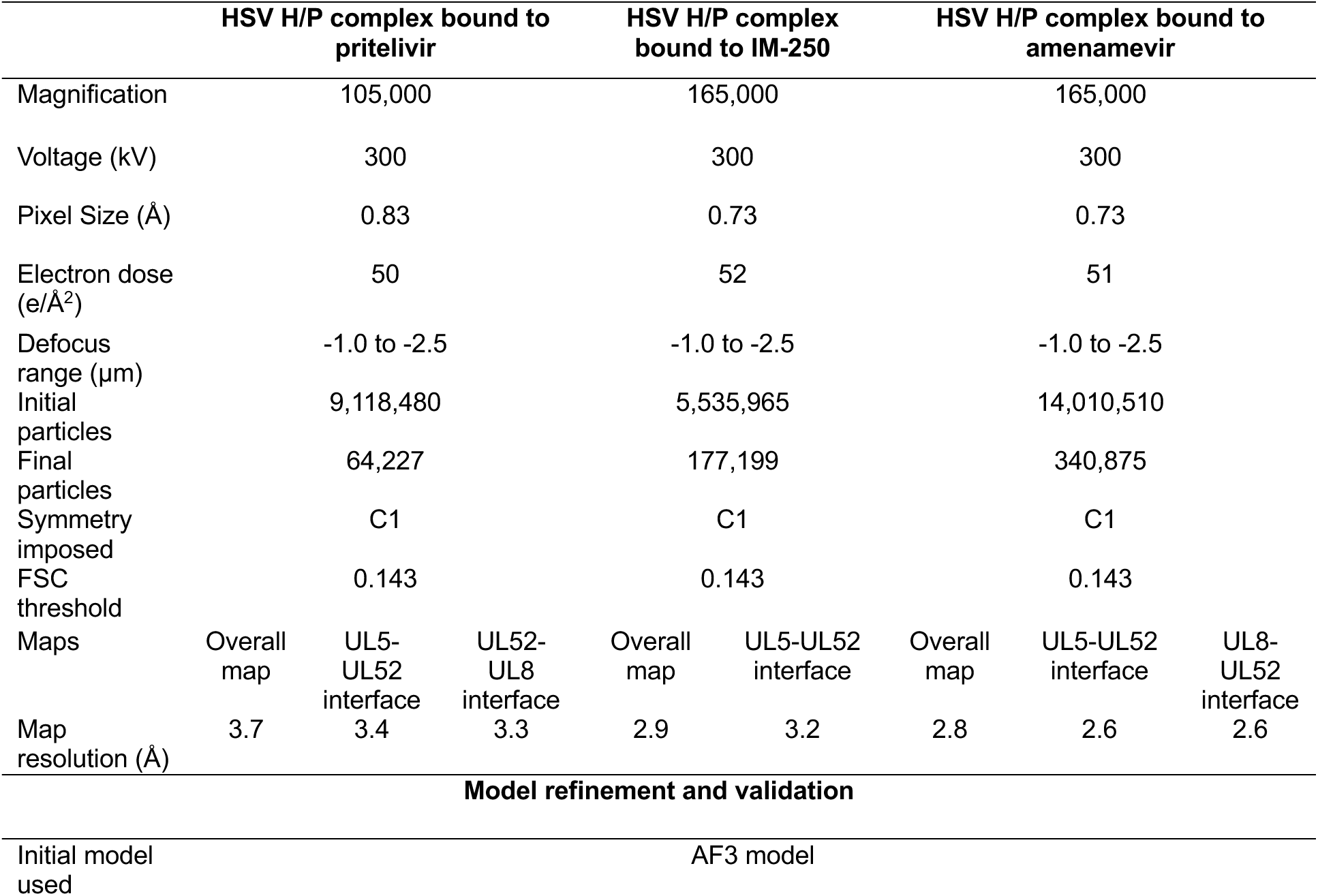
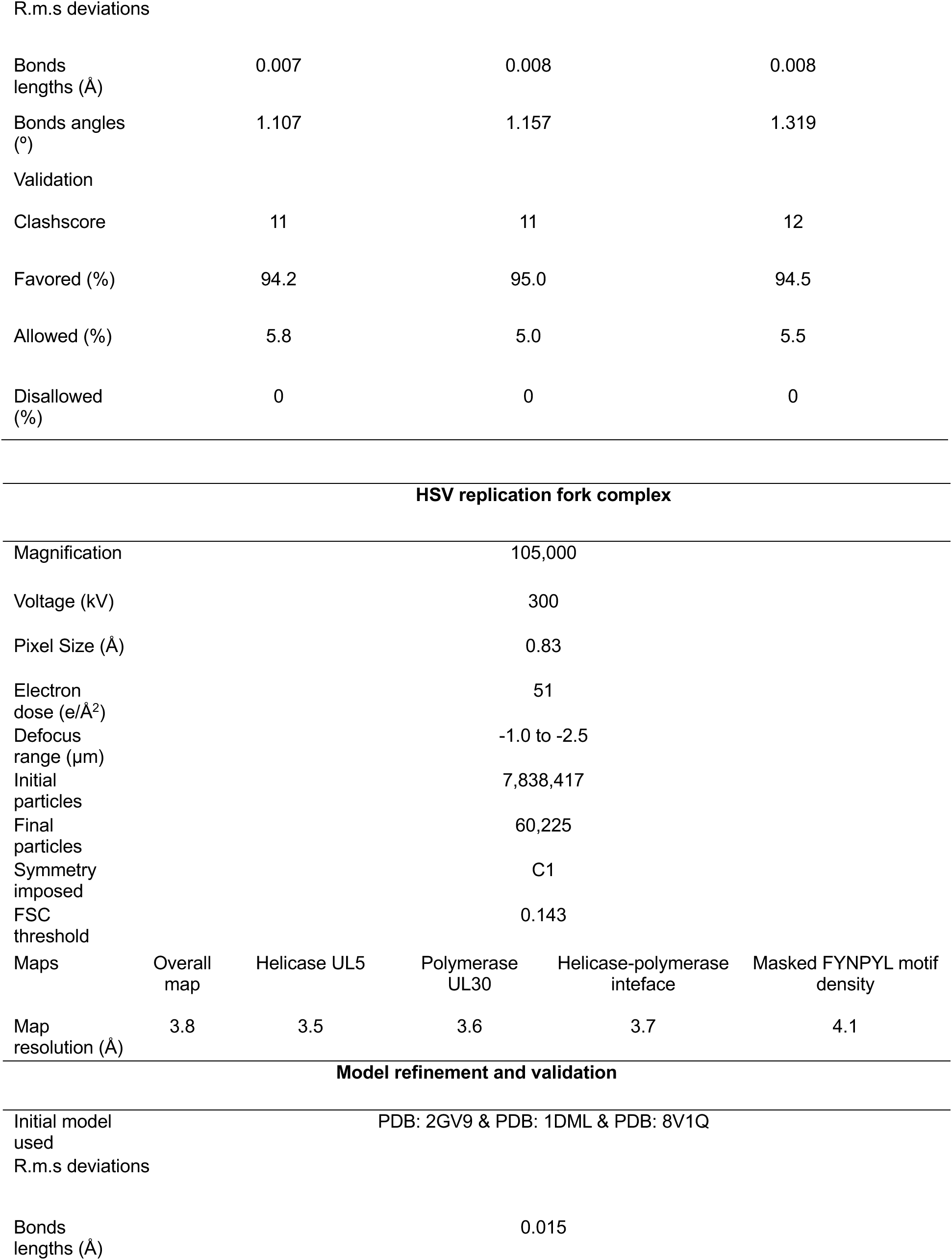
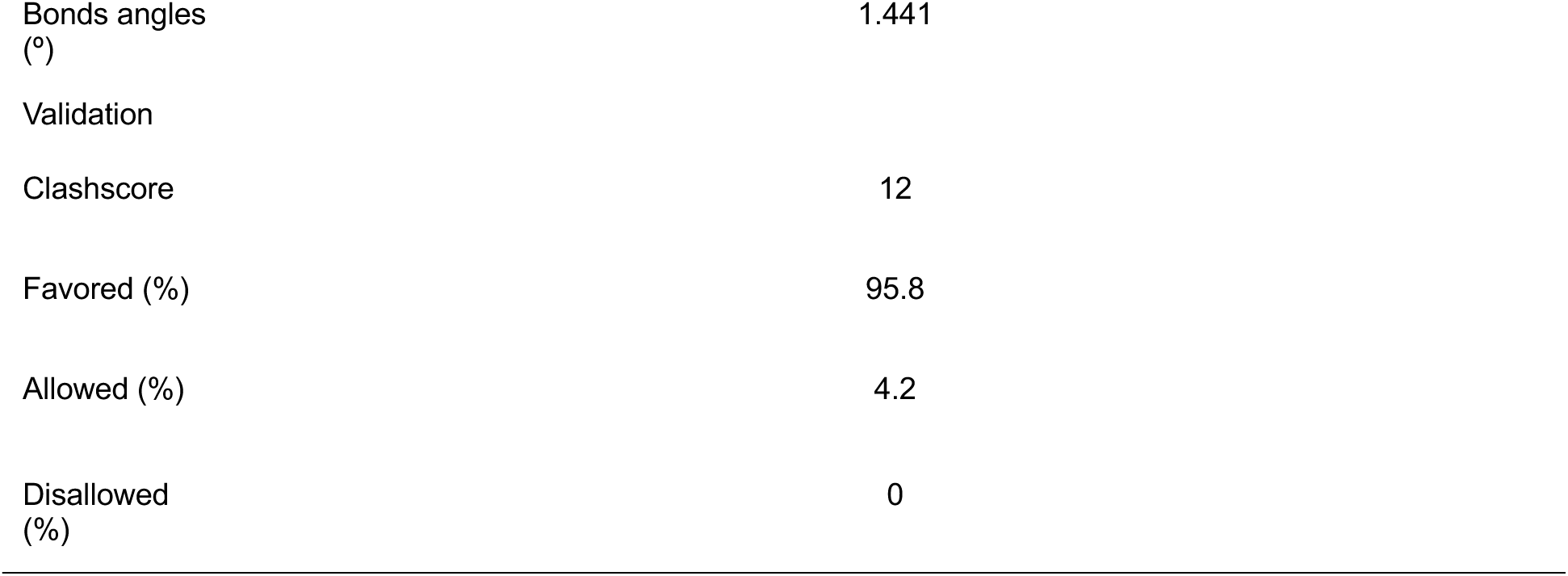
Cryo-EM data collection and image processing for drug-bound H/P complex structures, related to Figures 1 and S1.

|  | HSV H/P complex bound to pritelivir |  |  | HSV H/P complex bound to IM-250 |  | HSV H/P complex bound to amenamevir |  |  |
| --- | --- | --- | --- | --- | --- | --- | --- | --- |
| Magnification | 105,000 |  |  | 165,000 |  | 165,000 |  |  |
| Voltage (kV) | 300 |  |  | 300 |  | 300 |  |  |
| Pixel Size (Å) | 0.83 |  |  | 0.73 |  | 0.73 |  |  |
| Electron dose (e/Å<sup>2</sup>) | 50 |  |  | 52 |  | 51 |  |  |
| Defocus range (μm) | -1.0 to -2.5 |  |  | -1.0 to -2.5 |  | -1.0 to -2.5 |  |  |
| Initial particles | 9,118,480 |  |  | 5,535,965 |  | 14,010,510 |  |  |
| Final particles | 64,227 |  |  | 177,199 |  | 340,875 |  |  |
| Symmetry imposed | C1 |  |  | C1 |  | C1 |  |  |
| FSC threshold | 0.143 |  |  | 0.143 |  | 0.143 |  |  |
| Maps | Overall map | UL5-UL52 interface | UL52-UL8 interface | Overall map | UL5-UL52 interface | Overall map | UL5-UL52 interface | UL8-UL52 interface |
| Map resolution (Å) | 3.7 | 3.4 | 3.3 | 2.9 | 3.2 | 2.8 | 2.6 | 2.6 |
| **Model refinement and validation** |  |  |  |  |  |  |  |  |
| Initial model used | AF3 model |  |  |  |  |  |  |  |

R.m.s deviations
|  |  |  |  |
| --- | --- | --- | --- |
| Bonds lengths (Å) | 0.007 | 0.008 | 0.008 |
| Bonds angles (°) | 1.107 | 1.157 | 1.319 |
| Validation |  |  |  |
| Clashscore | 11 | 11 | 12 |
| Favored (%) | 94.2 | 95.0 | 94.5 |
| Allowed (%) | 5.8 | 5.0 | 5.5 |
| Disallowed (%) | 0 | 0 | 0 |

|  |  |  |  |  |  |
| --- | --- | --- | --- | --- | --- |
| Magnification | 105,000 |  |  |  |  |
| Voltage (kV) | 300 |  |  |  |  |
| Pixel Size (Å) | 0.83 |  |  |  |  |
| Electron dose (e/Å<sup>2</sup>) | 51 |  |  |  |  |
| Defocus range (µm) | -1.0 to -2.5 |  |  |  |  |
| Initial particles | 7,838,417 |  |  |  |  |
| Final particles | 60,225 |  |  |  |  |
| Symmetry imposed | C1 |  |  |  |  |
| FSC threshold | 0.143 |  |  |  |  |
| Maps | Overall map | Helicase UL5 | Polymerase UL30 | Helicase-polymerase interface | Masked FYNPYL motif density |
| Map resolution (Å) | 3.8 | 3.5 | 3.6 | 3.7 | 4.1 |

Model refinement and validation
|  |  |
| --- | --- |
| Initial model used | PDB: 2GV9 & PDB: 1DML & PDB: 8V1Q |
| R.m.s deviations |  |
| Bonds lengths (Å) | 0.015 |

|  |  |
| --- | --- |
| Bonds angles<br>(°) | 1.441 |
| Validation |  |
| Clashscore | 12 |
| Favored (%) | 95.8 |
| Allowed (%) | 4.2 |
| Disallowed<br>(%) | 0 |

**Table S4.** Summary of previously published mutational analyses.

| Subunit | Mutant | Location | Effects | Reference |
| --- | --- | --- | --- | --- |
| UL52 | C1023A<br>C1028A | ZnF zinc-coordinating residues | Near complete loss of helicase and primase activity of purified UL5–UL52 subcomplex. Mutant UL52 could not complement UL52-null virus, indicating loss of essential function. | Biswas et al.<sup>43</sup><br>Chen et al.<sup>44</sup> |
| UL52 | D628A<br>D630A | Primase active site residues involved in metal coordination | Decrease primase activity, no effect on helicase activity. | Dracheva et al.<sup>13</sup><br>Klinedinst et al.<sup>42</sup> |
| UL5 | D249A<br>E250A | ATPase active site residues involved in metal coordination | Mutations profoundly decrease DNA-dependent ATPase activity. Substitutions abolish DNA replication in cells. | Zhu et al.<sup>10</sup><br>Graves-Woodward et al.<sup>41</sup> |
| UL5 | K103A<br>R345K | ATPase active site residues involved in phosphate coordination | Substitutions of residues abolish DNA replication in cells. | Zhu et al.<sup>10</sup> |
| UL30 | FYNPYL motif mutant to poly A<sub>6</sub> | preN motif residues interacting with UL5 | No effects on polymerase DNA synthesis activity but impairs viral replication in cells. Severely impairs replication in sensory ganglia and latency establishment. | Terrell et al.<sup>18</sup><br>Terrell et al.<sup>19</sup> |

**Table S5.**
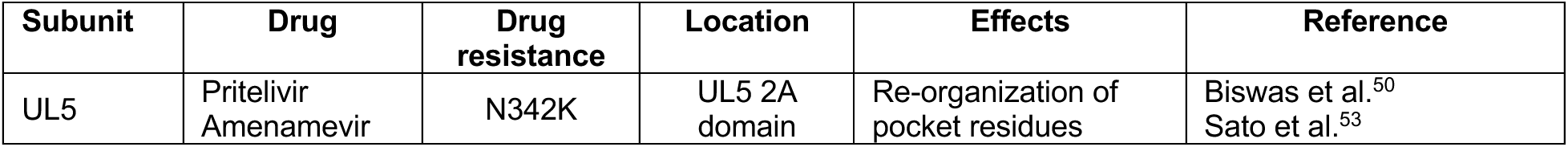
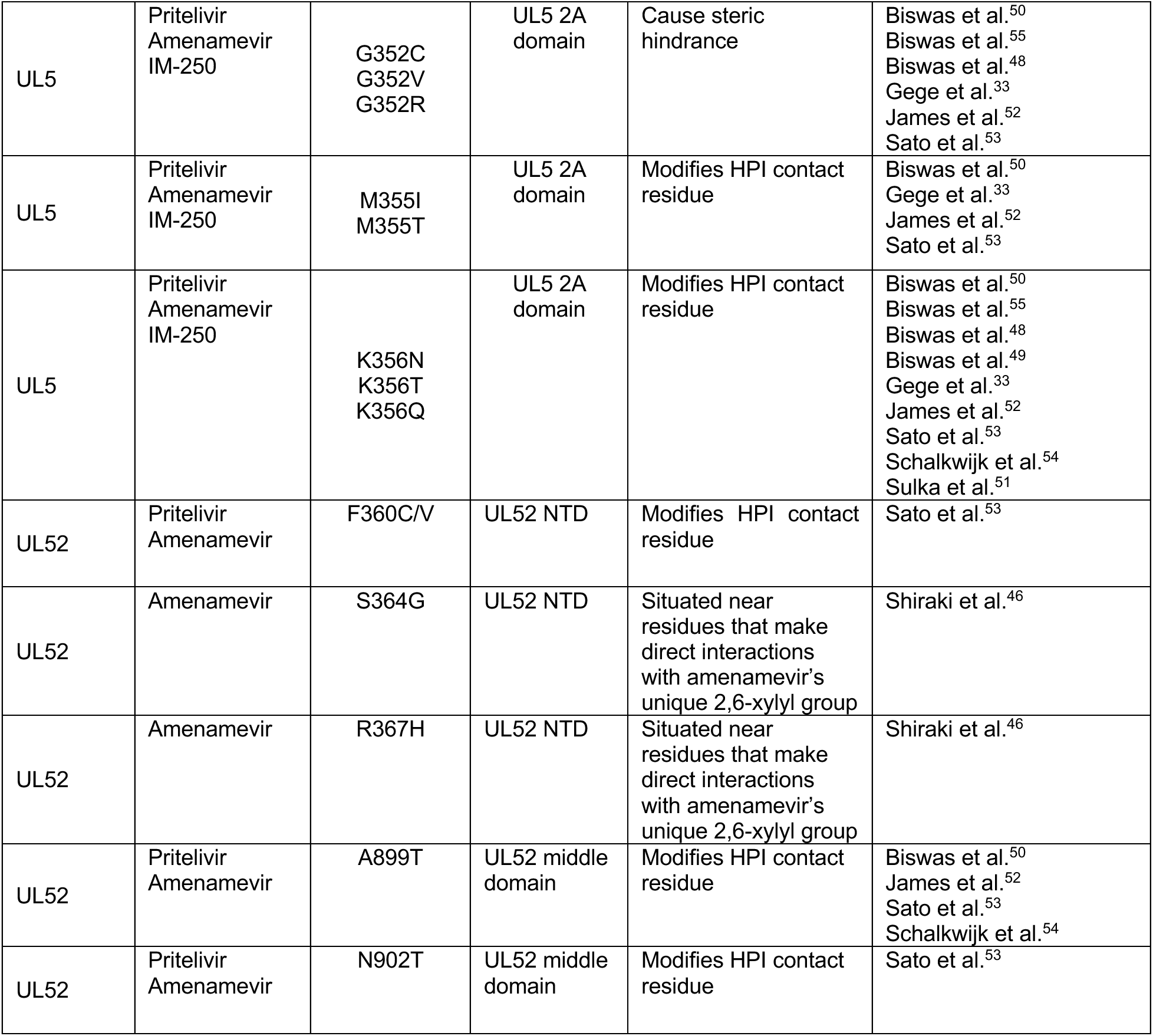
Summary of previously published drug-resistant mutation profiles.

